# Double-strand breaks induce inverted duplication chromosome rearrangements by a DNA polymerase δ and Rad51-dependent mechanism

**DOI:** 10.1101/2023.01.24.525421

**Authors:** Amr Al-Zain, Mattie R. Nester, Lorraine S. Symington

**Author notes:** To whom correspondence should be addressed: Lorraine S. Symington.

## Abstract

Inverted duplications, also known as foldback inversions, are commonly observed in cancers and are the major class of chromosome rearrangement recovered from yeast cells lacking Mre11 nuclease. Foldback priming at naturally occurring inverted repeats is one mechanism proposed for the generation of inverted duplications. However, the initiating lesion for these events and the mechanism by which they form has not been fully elucidated. Here, we show that a DNA double-strand break (DSB) induced near natural short, inverted repeats drives high frequency inverted duplication in Sae2 and Mre11-deficient cells. We find that DNA polymerase δ proof-reading activity acts non-redundantly with Rad1 nuclease to remove heterologous tails formed during foldback annealing. Additionally, Pol32 is required for the generation of inverted duplications, suggesting that Pol δ catalyzes fill-in synthesis primed from the foldback to create a hairpin-capped chromosome that is subsequently replicated to form a dicentric isochromosome. Stabilization of the dicentric chromosome after breakage involves telomere capture by non-reciprocal translocation mediated by repeat sequences and requires Rad51.

## INTRODUCTION

Most human cancer cells exhibit genomic instability, ranging from elevated mutation rates to gross chromosome rearrangements and aneuploidy. Gross chromosome rearrangements (herein called GCRs) include translocations, deletions, inversions and amplifications. The prevailing view is that GCRs are generated through error-prone processing of damaged chromosomes ^1^. Although the nature of the initiating lesions for GCRs is unknown, much of the genetic evidence from yeast and human cells implicates DNA double-strand breaks (DSBs) formed directly or indirectly by problems during DNA replication ^2^.

GCRs have been studied extensively in yeast using genetic assays developed by the Kolodner group ^3, 4^. The classical GCR assay detects the spontaneous loss of two counter-selectable genes, *CAN1* and *URA3*, that are present on the left arm of chromosome V (Chr V-L), telomeric to the last essential gene. Using this assay, the rates of GCRs in various mutant genetic backgrounds as well as the types of rearrangements have been well characterized. In wild-type (WT) cells, the most common type of rearrangement is telomere addition, in which terminal Chr V-L loss is accompanied by *de novo* addition of sequences telomeric to the break site ^2, 3^. Other rearrangements include interstitial deletions, non-reciprocal translocations and inverted duplications ^1^.

Inverted duplications, also called foldback inversions or palindromic duplications, are thought to arise from a DSB intermediate ^5–8^. There are several proposed models for the formation of inverted duplications initiated by DSBs and these can be broadly categorized as replicative or non-replicative. Non-replicative inverted duplications arise through fusion of sister-chromatids after replication of a chromosome that has sustained a DSB or telomere attrition during G1 phase ^9–11^. Sister-chromatid fusions occur mainly by microhomology-mediated end joining ^12–14^, but have also been reported to occur by single-strand annealing (SSA) involving long inverted repeats ^15^. Telomere-telomere fusions, in contrast, are largely NHEJ-dependent ^14, 16–18^. Replicative inverted duplications are thought to occur via intra-strand annealing between short, inverted repeats exposed by end resection^19–21^. Subsequent fill-in synthesis and ligation create a hairpin-capped chromosome. It has been suggested that during the next cell cycle, the replisome loops at the hairpin and replicates back to the end of the chromosome. Both sister-chromatid fusions and replication through hairpin-capped chromosomes create dicentric isochromosomes.

Dicentric chromosomes are unstable as they can be broken during cytokinesis when they are pulled apart by each daughter cell and form a bridge. Asymmetric breakage results in an inverted duplication, the degree of which depends on the location of the break. In budding yeast, breakage occurs either at the center of telomere-telomere fusions or within a 25-30 kb region near the centromere of dicentrics without telomere fusions ^22, 23^. Broken dicentrics can then undergo sister chromatid fusion, initiating a breakage-fusion-bridge cycle, or be stabilized by acquisition of a telomere either by *de novo* telomere addition or by BIR through a repeat sequence ^24, 25^. Alternatively, a dicentric can be stabilized by loss of one of its centromeres ^24, 26^.

The frequency of inverted duplications detected using the spontaneous GCR assay is elevated in *sae2*Δ mutants as well as cells defective for Mre11 nuclease activity ^5–7^. At the center of those inverted duplications are naturally occurring 3-11 bp long inverted repeats (IRs) ^5–7^, suggesting that they form via intra-strand foldback annealing between the IRs. Mre11 (as part of the Mre11-Rad50-Xrs2 complex) has endonuclease activity that is stimulated by Sae2 and is proposed to cleave hairpin-capped ends ^27–31^. Mre11-Sae2 are thus thought to prevent inverted duplications by resolving the hairpin-capped chromosome intermediate^5–8, 28, 32^.

Studies have shown that a DSB near long artificially integrated palindromes or quasi palindromes (40 bp or longer) can lead to inverted duplications through a hairpin-capped intermediate ^15, 33, 34^. DSBs near short IRs have also been shown to simulate the formation of inverted duplications ^7, 8^. However, the specific effect of the DSB on the frequency and the mechanism of GCRs has not been systematically studied. In this study, we monitored the repair outcome of a CRISPR/Cas9-induced DSB near a naturally occurring IR. Our data show that inverted duplications occur at a surprisingly high frequency in cells deficient for the Mre11 nuclease activity. Similar to previously proposed models, the inverted duplications occur through intra-strand foldback annealing at resected inverted repeats to form a hairpin-capped chromosome that is a precursor to dicentric isochromosomes. We identify two roles for DNA polymerase δ in the generation of foldback inversions. First, the proof-reading activity is required for removal of heterologous tails formed during foldback annealing, and second, the Pol δ processivity subunit Pol32 is important for fill-in synthesis to generate a hairpin-capped chromosome. We show that stabilization of dicentric chromosomes after breakage occurs by Rad51-dependent recombination between repeat sequences.

## RESULTS

### Cells lacking Mre11 endonuclease activity exhibit increased survival to a DSB induced near a natural inverted repeat

To determine whether a targeted DSB induces inverted duplications, particularly in the context of Mre11 nuclease deficiency, we used CRISPR/Cas9 ^35^ to create a DSB near a naturally occurring IR on Chr V-L. The IR chosen was observed to be at the center of four spontaneous inverted duplications previously recovered from *sae2*Δ cells ^6^ (Figure 1A). It is an imperfect repeat of 11 bp with two mismatches, separated by 6 bp, within the *CAN1* gene. The gRNA (hereafter referred to as gRNA-17) was designed to target a sequence 17-bp telomeric to the IR, thus creating a DSB 20 bp from the IR as Cas9 cuts 3 bp from the target sequence 3′ end. Sequences telomeric to the DSB are non-essential for cell viability, permitting recovery of a variety of GCRs. The gRNA-17 expression cassette was stably integrated into the genome.

**Figure 1.**
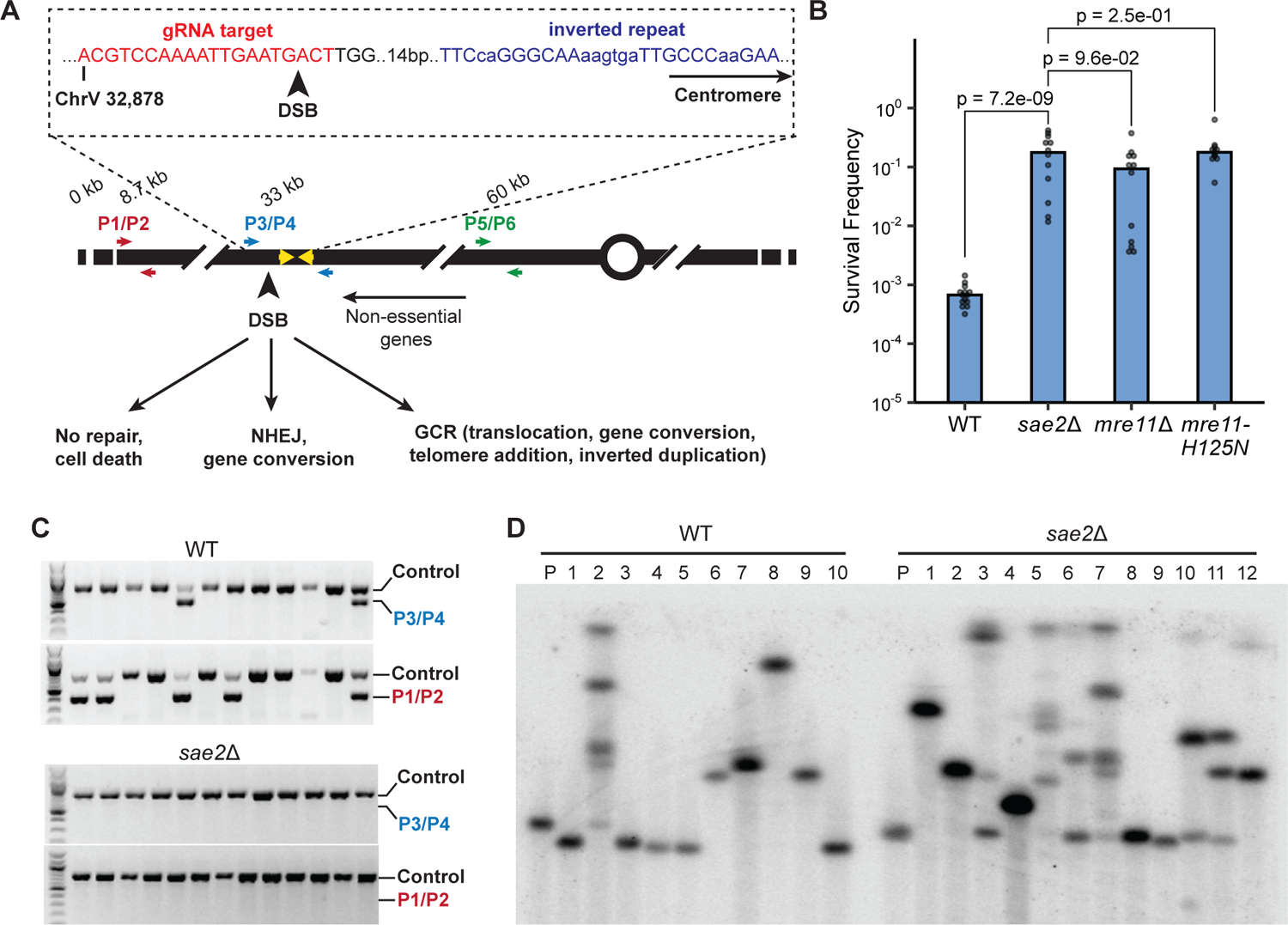
Cells lacking Mre11 endonuclease activity exhibit increased survival to a DSB induced near a natural inverted repeat. A. Top: Sequence of the gRNA target (red font) located telomere proximal to the targeted IR (blue font). The mismatches within the 11 bp imperfect repeats and the 6 bp spacer are shown in lower case. Bottom: Schematic of the left arm of Chr V showing the location of the DSB and inverted repeats (yellow arrows). Primer pairs to detect retention of terminal sequence, NHEJ events, and copy number increase are shown in red, blue and green, respectively. B. Survival frequencies of the indicated strains in response to expression of Cas9 and gRNA-17 (see Methods). P values were determined using a two-tailed t test. C. Twelve independent clones from WT and *sae2*Δ cells were analyzed using P3/4 and P1/2 primers to detect NHEJ events or loss of Chr V terminal sequence. D PFGE of 10 WT clones that were pre-screened to eliminate NHEJ and homeologous conversion events, and 12 *sae2*Δ clones analyzed directly from β-estradiol plates. Shown is a Southern blot with a probe that hybridizes to *PCM1* on Chr V.

We initially used the galactose-inducible *GAL1* promoter to drive expression of Cas9 but observed a high level of leaky expression when cells were grown under non-inducing conditions, as measured by inactivation of *CAN1* in the presence of gRNA-17 (Figure S1A). To reduce background cleavage by Cas9, we switched to an estrogen-inducible LexA hybrid transcription factor fusion to drive expression of Cas9 under the control of the *lexO* operator and a minimal *P_CYC1_* promoter ^36^ (Figure S1B). However, we still observed a high *CAN1* mutation rate under non-inducing conditions (Figure S1A). To reduce background cleavage by Cas9, we fused an ER domain to Cas9 (*lexO-Cas9-ER*). Using this system, Cas9 expression as well as nuclear localization is dependent on the addition of β-estradiol to the medium. A single cassette containing the transcription factor and *lexO-Cas9-ER* was stably integrated into the genome. Due to the presence of two inverted ER domains in the construct, there is a possibility for spontaneous recombination between them, which would disrupt the LexA-ER-AD transcription factor and eliminate Cas9 expression. Although these events were expected to be rare, they represent about 20% of the total number of colonies that grow on inducing medium in wild-type (WT) cells and were filtered out for subsequent analysis.

Logarithmically growing haploid cells were plated on media +/-β-estradiol and cell survival was determined by the ratio of colonies that grew on β-estradiol-containing medium versus medium lacking β-estradiol. Since Cas9 is constitutively expressed when cells are plated on medium containing β-estradiol, and there is no repair template for homologous recombination, cells can only grow if they lost the gRNA target sequence. Therefore, repair in surviving cells is likely to be inherently mutagenic. The gRNA target sequence can be lost by indels via NHEJ or larger scale sequence loss (Figure 1A). The survival frequency of WT cells was around 0.05% (Figure 1B), an order of magnitude lower than the survival of cells to DSBs generated by HO or I-SceI endonucleases ^37, 38^, suggesting that NHEJ is ineffective in repair of the Cas9-induced DSB. Remarkably, survival of *sae2*Δ cells (11.5%, ± 11.2) was ∼200-fold higher than that of WT cells (0.05%, ± 0.02%) (Figure 1B). The survival frequencies of *sae2*Δ*, mre11*Δ and *mre11-H125N* (deficient for the nuclease activity of Mre11) mutants were similar, suggesting more efficient stabilization of the broken chromosome than in WT cells.

The different potential repair outcomes are mutagenic NHEJ or chromosome rearrangements. To distinguish between these possibilities, we used PCR to detect the presence of 250 bp on either side of the break (Figure 1A, primer pair P3/P4). A PCR product indicates repair of the break without extensive loss of sequences, likely due to inaccurate NHEJ. PCR analysis of several individual clones derived from WT cells revealed that about 16% of survivors retained sequences surrounding the cut site (Figure 1C). Sequencing of P3/P4 PCR products (hereafter referred to as the cut site bands) revealed the presence of indels or base substitutions at the gRNA target site clustered in the first two nucleotides, or short deletions, indicative of NHEJ (Figure S2A). One possible explanation for the inefficiency of NHEJ is that Cas9-generated DSBs have mostly blunt ends ^35, 39^, which are poorly ligated by the NHEJ machinery in budding yeast ^40, 41^. *In vivo* studies imply that the ends generated by Cas9 can have 1-2 nt overhangs since the repair products often exhibit templated insertions, but this is influenced by sequence context of the DSB ^42–44^. Although *SAE2* deletion has been shown to increase the frequency of NHEJ ^45^, very few of the *sae2*Δ survivors had a cut site PCR product (Figure 1C). The significant increase in the survival frequency of those cells therefore reflects increased channeling to a repair pathway that leads to sequence loss.

To further characterize the mutagenic repair outcomes, we screened survivors by PCR using sub-telomeric primers (P1/P2 in Figure 1A) to determine whether the terminal sequences of Chr V-L were retained. Clones in which the terminal sequences are retained, but not the cut site sequences (three shown in Figure 1C) may be indicative of interstitial deletions. Alternatively, sequences that anneal to the cut site primer could be altered without otherwise significant sequence loss. To distinguish between the two outcomes, we used primers 200 bp further upstream and downstream of the cut site primers to screen survivors that only yielded the terminal fragment PCR (P7/P8, Figure S2B). P7/P8 gave a product in all eleven tested WT survivors that retained Chr V-L terminal PCR product but not the P3/P4 PCR product (Figure S2C). Sequencing of those PCR products indicates that repair occurred by gene conversion using the *LYP1* gene on Chr XIV as a template (Figure S2D). *LYP1* encodes a lysine permease that has only 61.6% sequence homology to *CAN1* and was not expected to template homology-directed repair at a detectable frequency ^46–48^. In a previous study, rare rearrangements between the highly diverged *CAN1* and *LYP1* genes were detected only in the absence of Sgs1 helicase ^49^, whereas here they were detected in 40% of the WT clones (Figure 1C and 2C).

**Figure 2.**
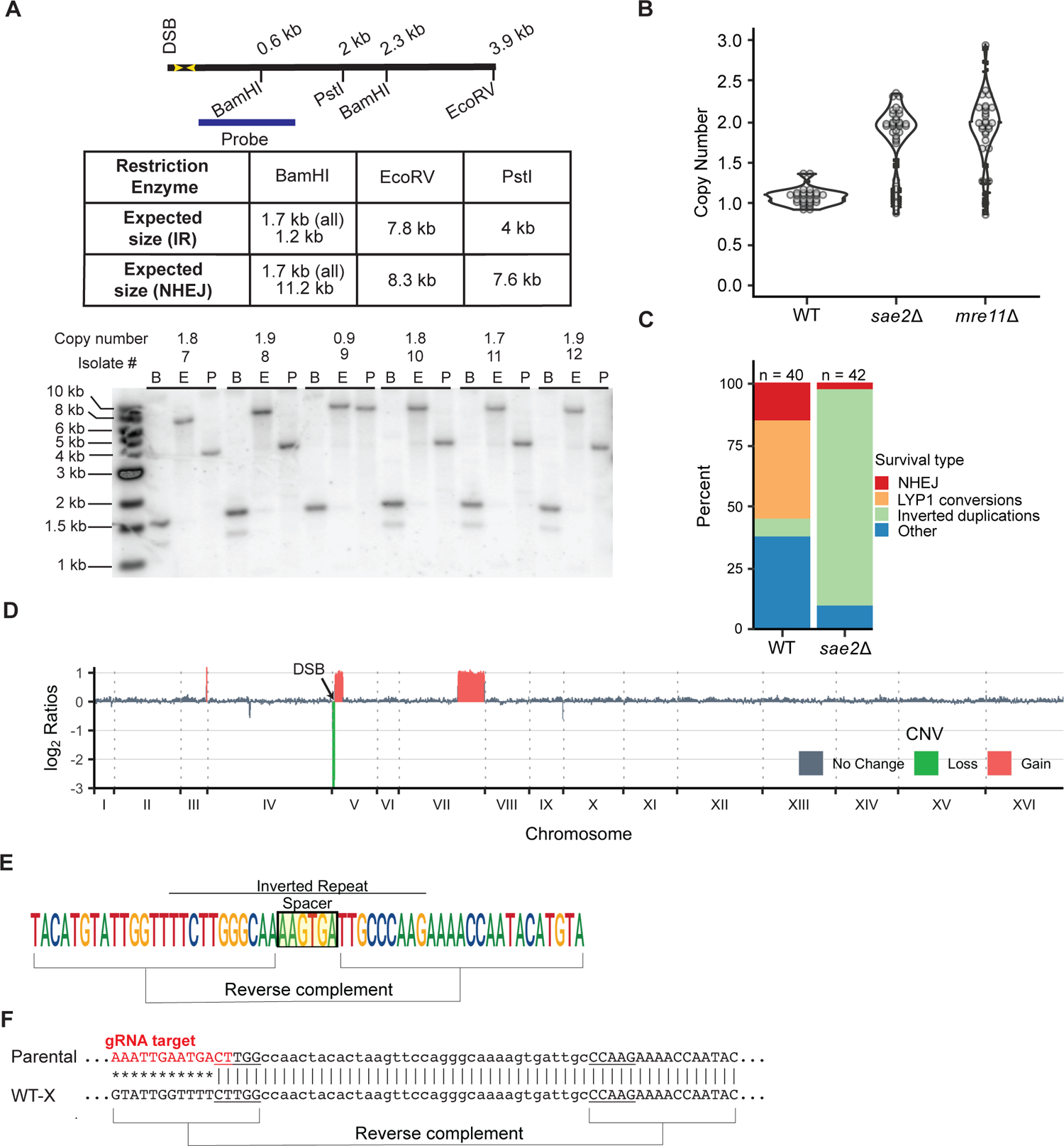
Mre11 endonuclease activity suppresses inverted duplications. A. The schematic indicates the distance of restriction endonuclease sites from the IR and expected sizes of fragments from inverted duplication clones. Genomic blot of DNA isolated from 6 *sae2*Δ clones digested with BamHI, EcoRV or PstI. The copy number of sequences detected by P5/6 primers, as determined by qPCR, are shown above each clone analyzed. B. Independent clones derived from WT, *sae2*Δ or *mre11*Δ strains were analyzed by qPCR (P5/6 primers) to detect increased copy number of sequence between the targeted IR and centromere (see Methods). C. Distribution of survivor types in WT and *sae2*Δ cells plotted as percent of total events analyzed. D. WGS of a *sae2*Δ clone showing sequence loss centromere distal to the targeted DSB, sequence duplication centromere proximal to the DSB, and a secondary event involving partial duplication of Chr VII. E. The *sae2*Δ clones analyzed had all corrected the mismatches present in the imperfect IR to generate a perfect palindromic duplication. F. The center of inverted duplication detected in one wild-type clone. Top: the inverted repeat from which the inverted duplication initiated. Underline is the inverted repeat and in lower case the spacer between repeats. Bottom: the sequence at the center of the inverted repeated as detected by WGS; * indicates sequence lost centromere distal to the IR.

To determine the size of Chr V in WT and *sae2*Δ clones, we performed pulsed-field gel electrophoresis (PFGE) of intact chromosomes and probed for Chr V (Figure 1D). The WT clones for PFGE analysis were pre-screened by PCR using P3/4 and P7/8 primer sets to eliminate those resulting from repair by NHEJ or homeologous gene conversion, whereas the *sae2*Δ clones were not pre-screened. Compared to the parental strain, Chr V of WT survivors was of aberrant size. Half exhibited chromosome truncations (#1, 3, 4, 5 and 10 in Figure 1D), characteristic of telomere addition or interstitial deletions. The remainder exhibited chromosome expansions, likely due to translocations or inverted duplications. One clone exhibited multiple bands (#2 in Figure 1D), indicative of a heterogenous population of cells that have undergone different rearrangements. Most survivors from *sae2*Δ cells exhibited chromosome expansions, some of which have multiple bands that hybridize with the Chr V probe, reminiscent of *sae2*Δ inverted duplication clones recovered using the classical GCR assay ^6^.

### Mre11 endonuclease activity suppresses DSB-induced inverted duplications

To test whether the GCRs in the *sae2*Δ clones that survived DSB formation are due to inverted duplications, genomic DNA from twelve survivors was analyzed by restriction endonuclease digestion and Southern blotting (Figure 2A). Restriction digestion of DNA with an inverted duplication would result in bands that are twice the size of the distance of the restriction site from the inverted duplication center. Eleven of twelve clones analyzed had restriction fragments consistent with the presence of an inverted duplication (#7, 8, 10, 11 and 12 in Figure 2A). The remaining clone had repaired the DSB by NHEJ (#9 in Figure 2A). In addition, we used a real time PCR (qPCR) strategy to screen a larger number of clones for copy number change in a region centromeric to the cut site (primers P5/P6, Figure 1A) relative to a control locus in a different chromosome (*ADH1*). The majority of *sae2*Δ and *mre11*Δ survivors had an increased copy number compared with WT (Figure 2B), indicative of duplications and consistent with the restriction digestion analysis. The qPCR and Southern blot analyses revealed that most of the *sae2*Δ survivors have inverted duplications, whereas this class of GCR is rare in WT cells (Figure 2C).

To identify the DNA sequences at the centers of inverted duplications, we prepared whole genome deep sequencing libraries from inverted duplication clones recovered from *sae2*Δ cells. Illumina paired-end sequencing revealed copy number changes consistent with an inverted duplication initiated at the Cas9-induced DSB. Sequences telomeric to the DSB were lost, whereas a duplication was detected centromeric to the DSB (Figure 2D). Furthermore, most of the *sae2*Δ clones exhibited an additional duplication of sequences from another chromosome, these are described in detail below. We were able to obtain the sequence at the breakpoint for the inverted duplications using Comice, which is part of the Pyrus software suite ^5^, and by *de novo* assembly of discordant reads pairs of which one read maps to the vicinity of the DSB site. For all *sae2*Δ clones sequenced, the inverted duplication was centered on the natural IR targeted by gRNA-17 (hereafter referred to as the target IR), consistent with the IR driving inverted duplications (Figure 2E, Table 1). Interestingly the two mismatches in the 11 bp imperfect inverted repeat were corrected to match the centromere proximal repeat. This finding suggests that the 3’ end to be extended after forming a fold back undergoes polymerase proofreading or mismatch repair. Surprisingly, we found that two of three inverted duplications recovered from WT cells used a different 5 bp-long IR that is located only one bp from the DSB and has a spacer of 35 bp (Figure 2F), while the other had a complex rearrangement at its center.

**Table 1.** Target IR and repeat elements used for secondary rearrangements of inverted duplication events

| Genotype | IR used |  |  | Secondary event repeat element |  |  | Total |
| --- | --- | --- | --- | --- | --- | --- | --- |
|  | Target IR | Long spacer | Other IR | *Delta* | *PAU* | Other |  |
| *sae2Δ* | 10 | 0 | 0 | 7 | 3 | 0 | 10 |
| *sae2Δ rad51Δ* | 9 | 0 | 1 | 8 | 2 | 0 | 10 |
| *sae2Δ pol32Δ* | 5 | 0 | 1 | 4 | 1 | 1<sup>a</sup> | 6 |
| *sae2Δ rad1Δ* | 8 | 0 | 0 | 4 | 1 | 3<sup>b</sup> | 8 |
| *sae2Δ rad1Δ pol3-01* | 3 | 2 | 0 | 2 | 2 | 1<sup>c</sup> | 5 |
| WT | 0 | 2 | 1 | 3 | 0 | 0 | 3 |

<sup>a</sup> Isochromosome
<sup>b</sup> Two complex rearrangements and one telomere addition event
<sup>c</sup> Isochromosome

Overall, these data indicate that a DSB near an inverted repeat is sufficient to induce a high frequency of inverted duplications in *sae2*Δ and *mre11*Δ backgrounds. Furthermore, they suggest that mutagenic repair that leads to GCRs is strongly suppressed in WT cells, with the nuclease activity of Mre11 playing a major role within the context of a DSB near short, inverted repeats.

### Inverted repeats proximal to a DSB are necessary for generation of inverted duplications

To confirm that the target IR is important for the generation of inverted duplications, we scrambled the genomic sequence corresponding to the IR by CRISPR-Cas9-mediated gene editing leaving the sequence targeted by gRNA-17 intact (Figure S3A). Scrambling the IR reduced the survival of *sae2*Δ cells by about 10-fold (Figure 3A). Seven percent of the survivors had inverted duplications, compared to 88% of *sae2*Δ cells with the natural inverted repeat (Figures 3B and 3C). The inverted duplications in the strain with the scrambled IR initiated from other inverted repeats (Figure S3B). Interestingly, 34% of the survivors with the scrambled IR have cut site PCR products indicative of NHEJ (Figures 3C and S3C), consistent with previous observations that *sae2*Δ increases the frequency of NHEJ ^45, 50^. Overall, these results suggest that an IR proximal to a DSB is required for formation of inverted duplications, likely functioning by forming a hairpin-capped chromosome via a foldback mechanism.

**Figure 3.**
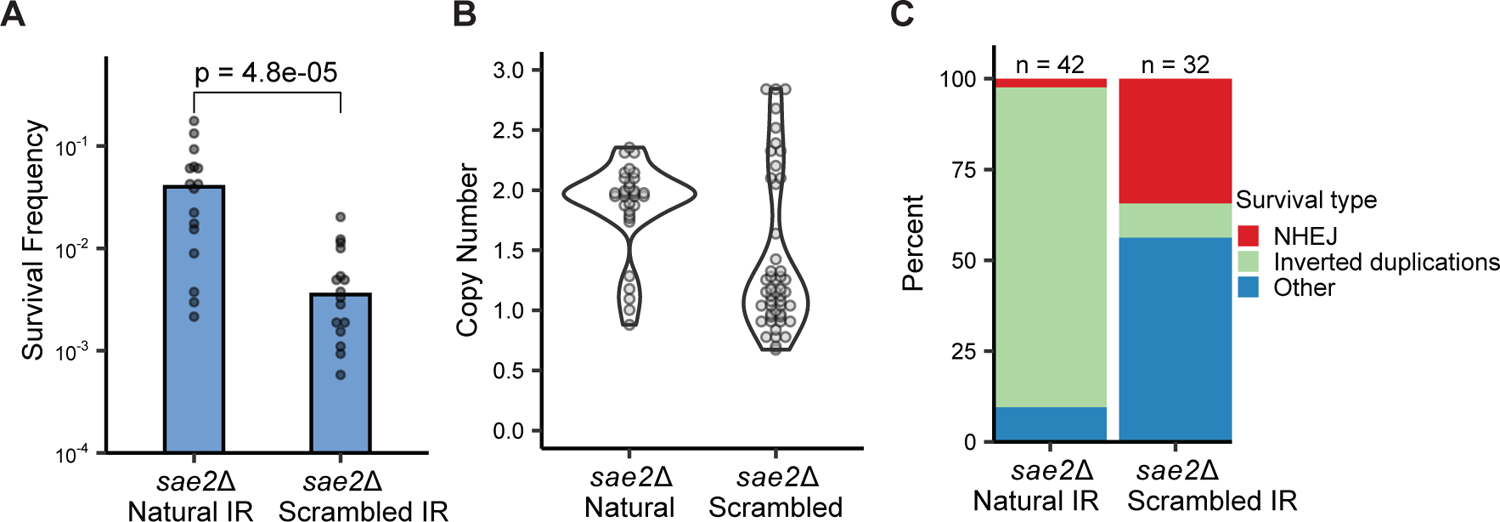
The targeted IR is required for inverted duplications. A. Survival frequencies of *sae2*Δ strains with the natural or scrambled IR in response to expression of Cas9 and gRNA-17. P value was determined using a two-tailed t test. B. Independent clones of the indicated genotype were analyzed by qPCR to detect increased copy number of sequence between the targeted IR and centromere. C. Distribution of survivor types in WT and *sae2*Δ cells plotted as percent of total events analyzed.

### The proof-reading activity of DNA Pol δ contributes to heterologous flap removal

The initiating DSB is located 20 bp from the IR; therefore, formation of a hairpin-capped chromosome would require cleavage of a 20 nucleotide (nt) long heterologous flap (Figure 4A). The Rad1-Rad10 complex has been shown to cleave 3′ heterologous flaps generated during recombination ^51, 52^. Heterologous flaps of 20 nt or less can also be removed by the 3′-5′ proof-reading exonuclease activity of DNA polymerase δ ^53^. Thus, we tested the requirement for Rad1 and DNA Pol δ in the formation of inverted duplications. Cells lacking *SAE2* and *RAD1* survive the DSB to a similar degree as *sae2*Δ cells (Figure 4B), and most of the survivors tested exhibited duplications (Figure 4C). This finding suggests that Rad1-Rad10 is dispensable for the formation of inverted duplications when the flap is 20 nt long, consistent with a previous report on its activity on HR substrates with flaps of a similar length ^53^. In contrast, *sae2*Δ *pol3-01* cells, which have a defect in the catalytic activity of the proof-reading domain of Pol δ ^54^, showed significantly lower survival than *sae2*Δ and *sae2*Δ *rad1*Δ mutants and reduced duplications in the survivors (Figure 4B, C). Furthermore, *sae2*Δ *pol3-01 rad1*Δ cells showed similar survival to *sae2*Δ *pol3-01*, suggesting a non-redundant role for proof-reading activity of Pol δ for inverted duplication formation. Consistent with this conclusion, 27% of clones from *sae2*Δ *pol3-01* survivors retained sequences surrounding the cut site, indicative of NHEJ (Figure S4A). By contrast, all the clones analyzed from *sae2*Δ *rad1*Δ cells had lost sequences surrounding the cut site. The inverted duplication defect in *sae2*Δ *pol3-01* is surprising, since it was previously reported that Rad1-Rad10 can substitute for the proof-reading activity of Pol δ during flap removal when the flaps are <20 nt long ^53^.

**Figure 4.**
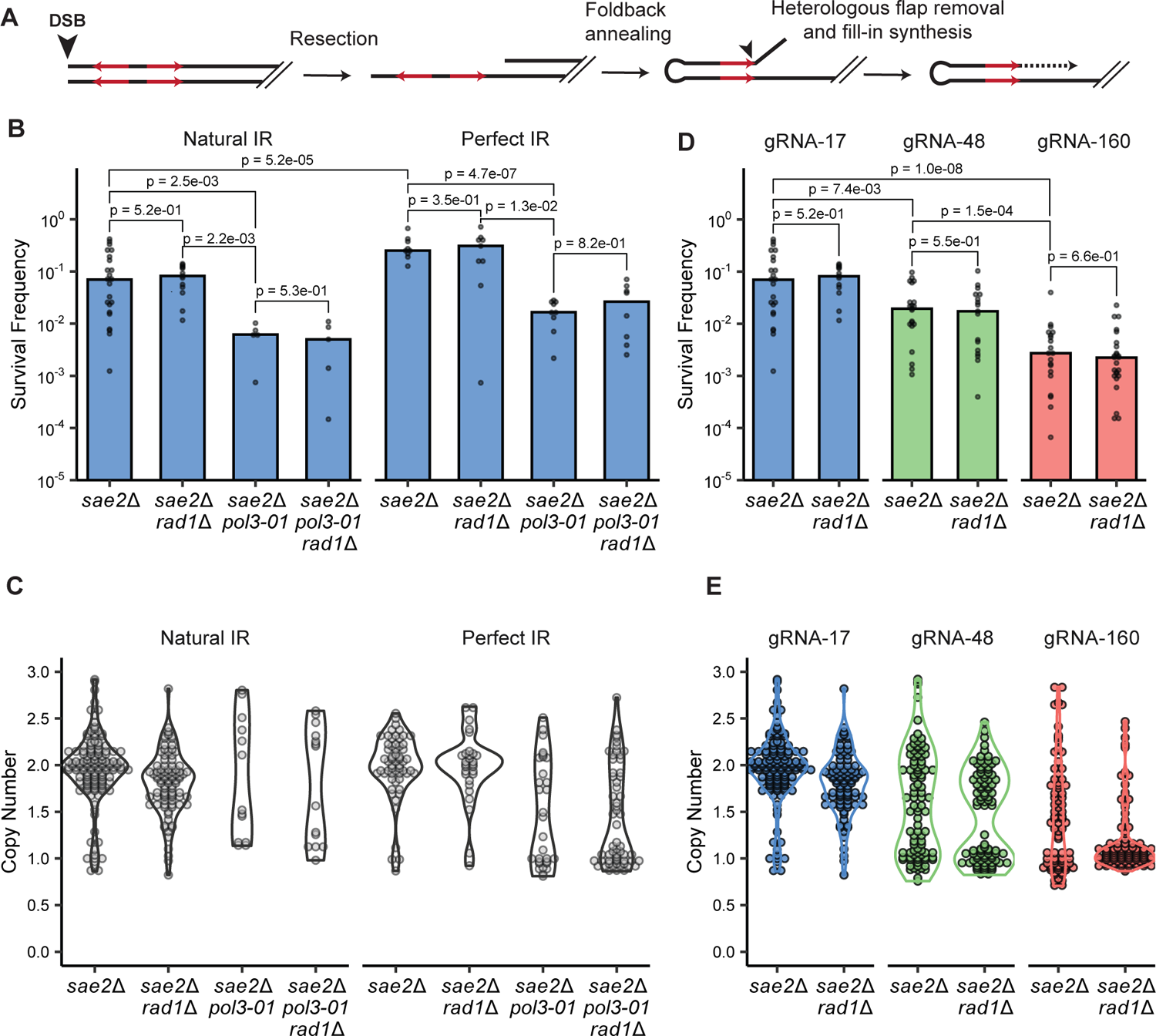
The DNA Pol δ proofreading activity is required for inverted duplication. A. Schematic showing the need for heterologous flap removal prior to fill-in DNA synthesis to form inverted duplications. B. Survival frequencies of the indicated genotypes with either the natural or perfect IR in response to expression of Cas9 and gRNA-17. P values were determined using a two-tailed t test. C. Independent clones of the indicated genotypes were analyzed by qPCR to detect increased copy number of sequence between the targeted IR and centromere. D. Survival frequencies of *sae2*Δ and *rad1*Δ strains in response to expression of Cas9 and gRNA-17, gRNA-48 or gRNA-160. P values were determined using a two-tailed t test. E. Independent clones of the indicated genotypes were analyzed as in C.

Deep sequencing of four inverted duplication clones from the *sae2*Δ *pol3-01 rad1*Δ triple mutant showed that two used the target IR and the other two used the IR with a 35-bp spacer that is located closer to the DSB, as observed for two of the WT clones, and thus would create a heterologous flap of only 1-nt (Figure S4C, Table 1). By contrast, all eight of the *sae2*Δ *rad1*Δ inverted duplications sequenced were centered on the target IR. These data confirm the importance of Pol δ proofreading activity in removal of heterologous flaps.

The defect in forming inverted duplications in the *sae2*Δ *pol3-01* mutant could be a consequence of the mismatches present in the target IR. Following foldback annealing, there is a 2-bp mismatch 3 bp away from the end of the repeat. If heterologous flap cleavage occurs at the end of the repeat, the remaining mismatches may present an obstacle for fill-in synthesis if not corrected and could contribute to the requirement for Pol δ proofreading activity. Consistent with this idea, Pol δ proofreading activity removes mismatches close to the 5′ invading end during homology-directed repair ^46, 55^. The sequencing data indicate that these mismatches are corrected in all inverted duplication clones, including clones derived from the *sae2*Δ *pol3-01 rad1*Δ cells. To determine whether the reduced frequency of inverted duplications in *sae2*Δ *pol3-01* cells is due to a defect in mismatch correction rather than in heterologous flap cleavage, we modified the chromosomal sequence to perfect the targeted IR (Figure S4B). The survival frequency of *sae2*Δ cells with the perfect inverted repeat was slightly increased relative to the original inverted repeat but survival and inverted duplication were still highly dependent on Pol δ proofreading activity (Figure 4B, C). Thus, the *pol3-01* defect is due to its essential role in heterologous flap removal and correction of mismatches within the foldback likely occurs by the canonical mismatch repair pathway.

Increasing the distance between the DSB and the IR is expected to decrease the frequency of inverted duplications if removal of long, heterologous flaps is inefficient or if a long flap destabilizes base pairing between the repeats. We designed two additional gRNAs that target sequences that are 48 and 160 bp away from the inverted repeat (Figure S4D). As predicted, the survival frequency of *sae2*Δ cells decreased as the DSB distance from the IR was increased (Figure 4D) and fewer inverted duplications were recovered (Figure 4E). This finding supports the hypothesis that the inverted duplications observed initiate by foldback intra-strand annealing at the IR. Although survival of *sae2*Δ *rad1*Δ cells using gRNA-48 or gRNA-160 was not significantly different to the *sae2*Δ single mutant, there was a decrease in the number of inverted duplications in the *sae2*Δ *rad1*Δ cells using gRNA-160 (*p*=0.002) (Table S1). To determine whether events scored as inverted duplications had initiated at a different IR to the target IR, we used a PCR strategy to detect sequence located between the binding site for gRNA-17 and the target IR (see Methods for details). The fraction of inverted duplications centered on the target IR decreased as the distance of the DSB from the target IR increased, and this effect was accentuated in the *rad1*Δ derivative using gRNA-48 (*p*=0.013) (Table S1). Furthermore, there was a bias toward use of an IR centromeric to the target IR in the *sae2*Δ strain expressing gRNA-48, whereas most of the inverted duplications from the *sae2*Δ *rad1*Δ strain had used an alternate IR telomeric to the target IR (*p*=0.003) (Table S1). These data are consistent with a minor role for Rad1-Rad10 nuclease in processing long heterologous flaps.

### Evidence for a dicentric chromosome intermediate

We recovered two clones in which the inverted duplication spans the entirety of Chr V minus sequence telomeric to the DSB (a *sae2*Δ *rad1*Δ *pol3-01* clone, Figure 5A, and a *sae2*Δ *pol32*Δ clone, Figure 5B). PFGE analysis of the two clones shows each has a derivative chromosome V that is almost twice the size of the native chromosome (Figure 5C), indicative of an isochromosome. These events initiated at the IR indicating that they were not products of sister chromatid fusion. In both cases, we identified a breakpoint junction consistent with an interstitial deletion spanning one of the two copies of the Chr V centromere (Figure 5A and 5B, bottom). This result provides evidence for the model that foldback priming near a DSB leads to an inverted duplication that extends from the break to the other end of the chromosome resulting in creation of a dicentric chromosome.

**Figure 5.**
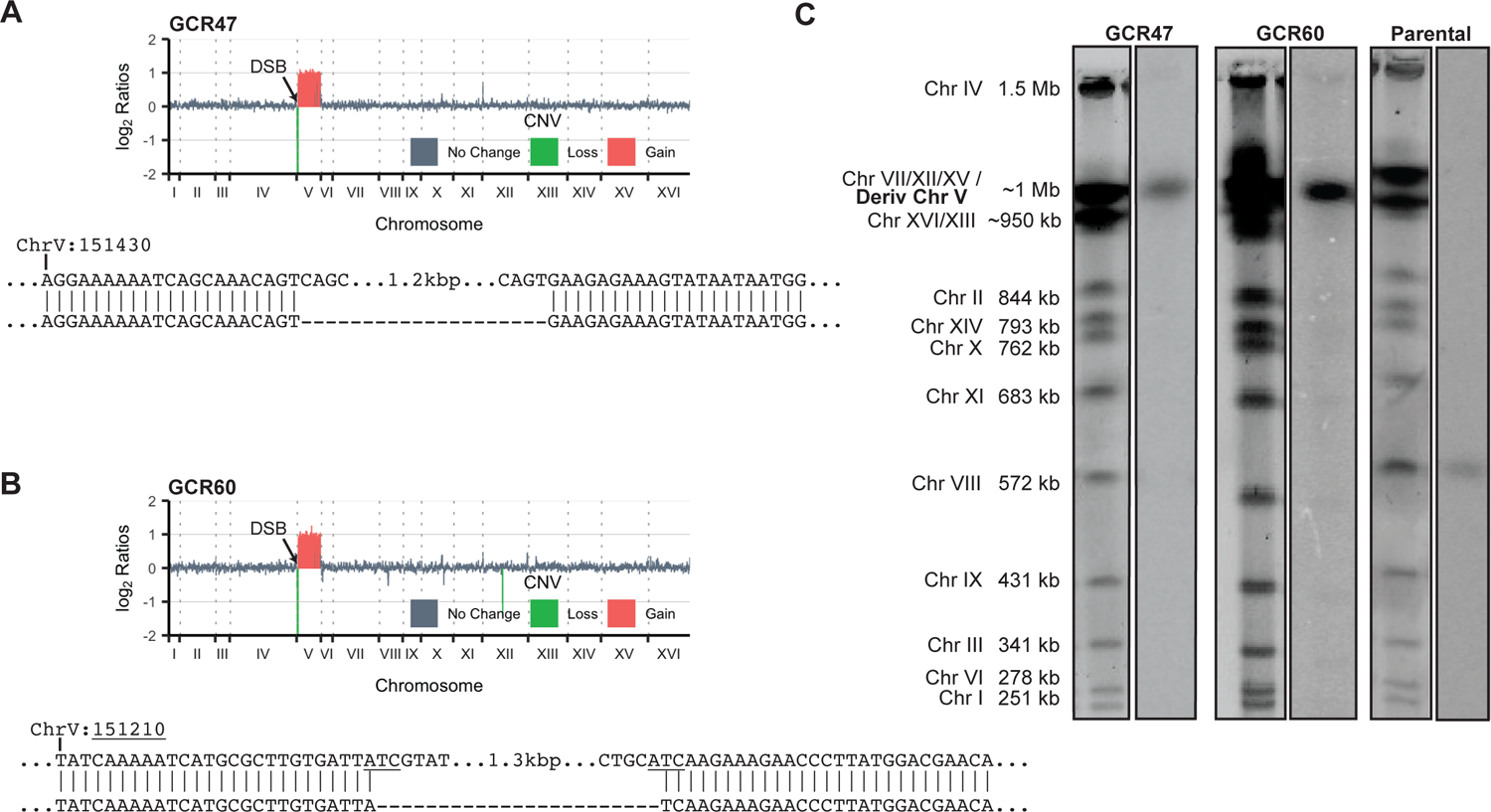
Inverted duplication extends the entirety of Chr V. A. and B. Top: Copy number analysis of two clones that exhibit completely duplication of Chr V centromeric to the targeted IR. Below: the sequence of junctions spanning a deletion of the *CEN5*. C. PFGE of the same two clones followed by Southern blotting using a probe against *PCM1* on Chr V.

### Rad51 and Pol32 are required for the formation of inverted duplications

For a hairpin-capped chromosome to form, gap filling DNA synthesis must occur to catch up with extensive resection. Consistent with this idea, Pol32, a non-essential subunit of the DNA Pol δ complex, is required for the formation of hairpin-capped chromosomes in RPA-depleted cells ^56^. If the inverted duplications form after foldback-annealing at the IR, then their formation should require Pol32. As expected, deletion of Pol32 in the *sae2*Δ background significantly decreased the frequency of survival to the Cas9-induced DSB (Fig 6A), and mostly eliminated the incidence of inverted duplications (Fig 6B). However, we found that most of the survivors recovered from *sae2*Δ *pol32*Δ cells had inverted the sequence between the ER domains within the Cas9 expression cassette, ablating Cas9 expression. To eliminate the possibility of this recombination event, we compared survival of *sae2*Δ *pol32*Δ cells expressing Cas9 with Cas9-ER. Survival was even lower in response to Cas9 without the ER domain than with Cas9-ER, consistent with an important role for Pol32 in formation of inverted duplications. This finding contrasts to a previous report using the spontaneous GCR assay, showing little effect of *POL32* deletion on the frequency of inverted duplications ^7^. Because Pol32 is also required for BIR ^57^, the reduced survival of *sae2*Δ *pol32*Δ cells could be due to its role in the secondary rearrangement necessary to stabilize a broken dicentric chromosome. However, the observation that inverted duplications are recovered from *sae2*Δ *rad51*Δ cells at a higher frequency than *sae2*Δ *pol32*Δ cells (see below) suggests that the *pol32*Δ defect is at an earlier step.

**Figure 6.**
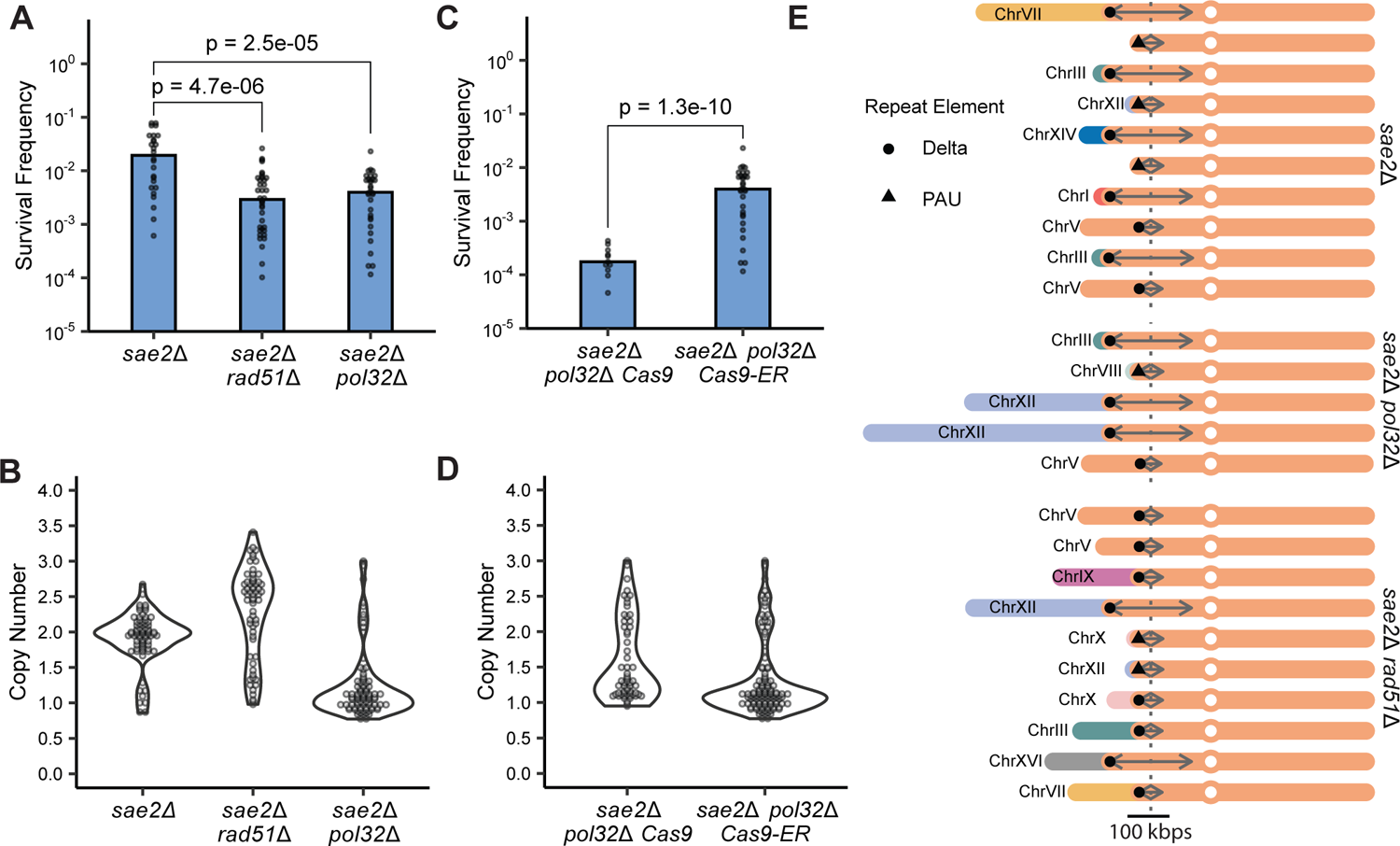
Rad51 and Pol32 contribute to inverted duplication formation. A. Survival frequencies of *sae2*Δ, *sae2*Δ *rad51*Δ and *sae2*Δ *pol32*Δ. P values were determined using a two-tailed t test. B. Independent clones of the indicated genotypes were analyzed by qPCR to detect increased copy between the targeted IR and centromere. C. Survival frequencies of *sae2*Δ *pol32*Δ strains wither either *CAS9* or *CAS9-ER*. P value was determined using a two-tailed t test. D. Copy number analysis by qPCR of the indicated strains was performed as B. E. Derivative chromosomes formed by rearrangements followed Cas9-induced DSB and inverted duplications as detected by WGS. Connectivity between the chromosomes was determined by analysis of the copy number change, by analysis for structural rearrangements using the Pyrus software suite and by de novo assembly of discordant reads. The dashed vertical line indicates the position of the DSB. Two-headed arrows denote the extend of the inverted duplication. Symbols denote the identity of the repeat sequence at the junction of the rearrangement.

Previous studies have shown that inverted duplications are frequently associated with homology-mediated secondary rearrangements ^5–7, 32^, presumably to stabilize a broken dicentric chromosome intermediate. Therefore, we anticipated survival would be reduced in *sae2*Δ *rad51*Δ cells. Although *sae2*Δ *rad51*Δ cells did indeed exhibit much lower survival to a DSB compared to *sae2*Δ, a majority of surviving cells were still able to form inverted duplications (Figure 6A, B).

### Secondary rearrangements associated with inverted duplications

To identify the extent of Chr V duplication and secondary recombination events, we analyzed the WGS data to detect copy number variation (CNV) genome wide (Figure 6E). Two main regions, at ∼62-65 kb and 135 kb, demarcate the end point of duplications, coinciding with the locations of delta elements and *PAU2*. *PAU2* is a member of the seripauperin gene family that are ∼350 bp in length and show a high degree of DNA sequence homology ^58^. There are approximately 21 *PAU* genes in the W303 *S. cerevisiae* genome, mostly located in sub-telomeric regions. In previous studies, the Ty1 element inserted at the *ura3* locus on Chr V-L was identified as a hotspot for secondary rearrangements associated with inverted duplications ^5–7^. Notably, the *ura3* locus in the W303 strain background used for this study lacks a Ty1 element and no inverted duplications terminated at this region of Chr V.

To determine the locations of translocations and whether repeat elements mediated them, we examined the structural variations present in *sae2*Δ clones using Comice ^5^. In *sae2*Δ, *sae2*Δ *rad51*Δ, and most *sae2*Δ *pol32*Δ inverted duplications, translocations were detected with the breakpoints aligning to either *PAU* genes or delta elements (Figure 6E, Table 1). In clones where there is no detectable CNV in a chromosome other than in Chr V, the translocations occurred with sub-telomeric sequences, which are difficult to map. Duplications followed by translocation with sub-telomeric regions (such as those mediated by *PAU* genes) result in derivative chromosome sizes that are not different from the parental Chr V. Therefore, a rearranged chromosome that migrates at the same distance as the parental Chr V by PFGE (Figure 1D) is not necessarily an indication that an inverted duplication has not occurred.

The secondary rearrangements that could be mapped in 5/6 *sae2*Δ *rad1*Δ and 4/4 *sae2*Δ *rad1*Δ *pol3-01* inverted duplications also appear to be mediated delta elements or *PAU2* (Figure S5A, Table 1). One *sae2*Δ *rad1*Δ secondary rearrangement was due to telomere addition. We also observed some complex rearrangements with evidence for quadruplications (Figures S5B, S5C and S5D). In each case, an inverted repeat marks the junction between a change in copy number. Such events could result from an additional round of foldback priming after breakage of a dicentric chromosome intermediate, as suggested previously ^6, 7^.

Three clones with inverted duplications from WT cells were sequenced and shown to have similar secondary rearrangements to the clones recovered from *sae2*Δ cells. In addition, we sequenced four WT GCR clones that lacked inverted duplications. In two of these, the rearrangements were due to microhomology mediated non-reciprocal translocations, another non-reciprocal translocation was mediated by 50 bp of sequence homology, and the fourth rearrangement was a Chr V truncation with *de novo* telomere addition (Figure S5E).

## DISCUSSION

In this work, we show how the DNA sequence context and genetic factors affect the outcome and frequency of DSB-induced inverted duplications. Previous studies have characterized inverted duplications that arise spontaneously ^5–7^. However, we find some differences in the genetic requirements and frequencies of inverted duplications when comparing spontaneous with DSB-induced events. This suggests that some spontaneous inverted duplications may arise through mechanisms that are different from those induced by DSBs and may not initiate from a DSB. We also find that a major determinant in channeling mutagenic repair is the sequence context of a DSB. In particular, a DSB near a short, naturally-occurring, inverted repeat is a potent inducer of foldback annealing, a precursor to inverted duplications. Mre11 and Sae2 play a significant role in resolving this intermediate and preventing foldback inversions.

The spontaneous GCR rate in *sae2*Δ cells is low (1.1 x 10^-9^) and is only about five-fold higher than in WT cells, with inverted duplications as the main class of GCR ^5, 6^. The increase in inverted duplications results from the loss of hairpin resolution activity of MRX-Sae2 ^28, 59, 60^. By contrast, the GCRs in WT cells are mainly telomere additions and inverted duplications are rare ^3, 5^. We found that ∼10% of *sae2*Δ cells can survive a DSB induced near a natural, short, inverted repeat, a >200-fold increase relative to WT cells, of which ∼90% were inverted duplications. This suggests that an initiating DSB is rate-limiting for the formation of inverted duplications in *sae2*Δ cells.

Inverted duplications in spontaneous GCR assays have been shown to contain short, inverted repeats at their centers that were derived from the original sequence ^5, 6^. This led to the hypothesis that these inverted duplications form by intra-strand foldback annealing of resected DNA at inverted repeats near the site of a DSB (Figure 7). It was recently shown that brief induction of a DSB in *sae2*Δ cells increased the rate of GCRs with inverted duplications. Those inverted duplications mainly used a perfect 15-bp IR, and the deletion of this hotspot did not reduce the GCR rate but changed the IRs used for inverted duplications ^7^. In congruence with the above studies, we found that introduction of a DSB near an 11-bp imperfect IR was both sufficient and necessary for the formation of inverted duplications in *sae2*Δ cells. Analysis of ten inverted duplication *sae2*Δ isolates by whole genome sequencing confirmed that the centers of the duplications lie at the targeted IR.

**Figure 7.**
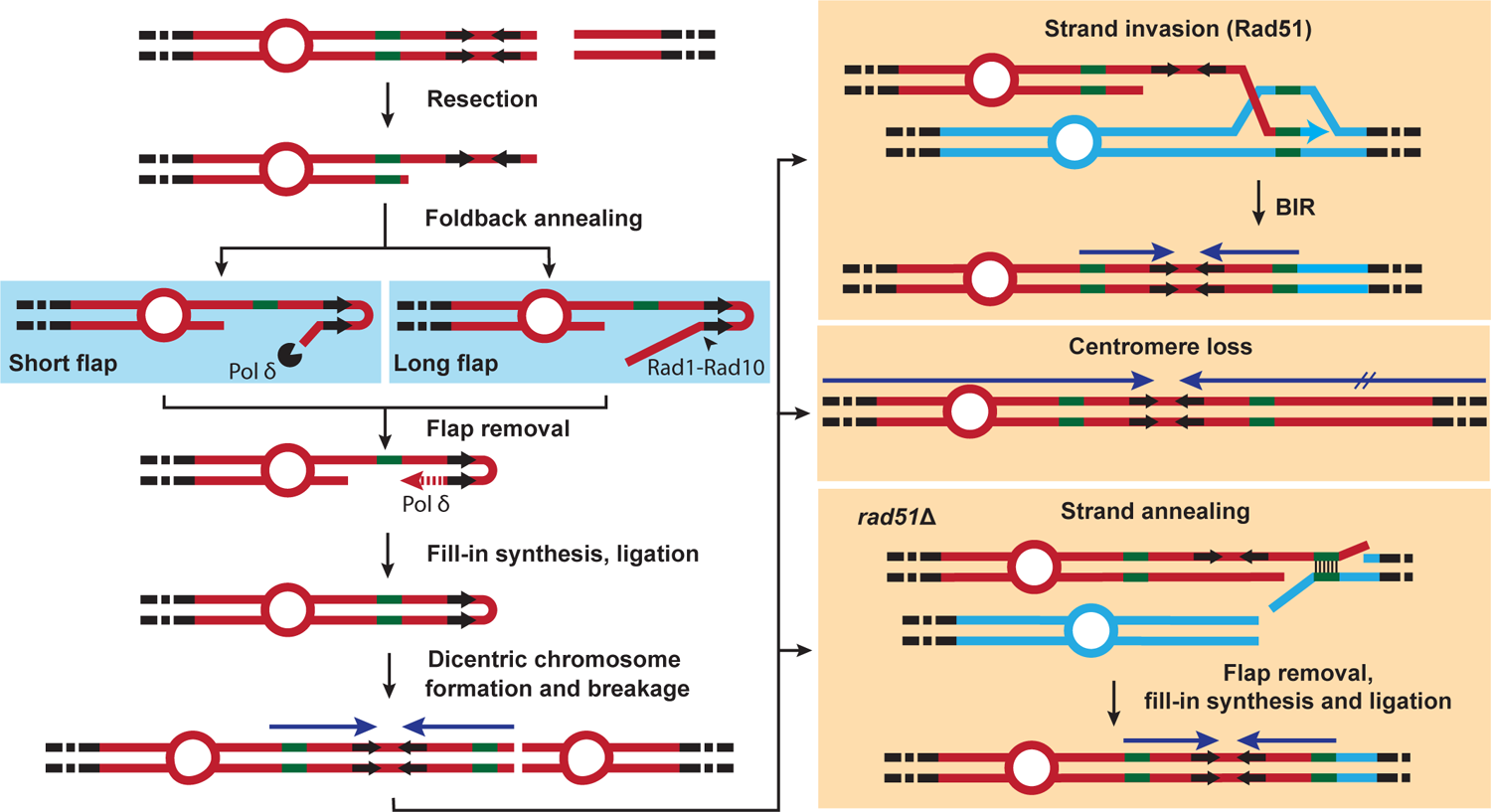
Model for inverted duplication formation. After DSB and resection, an exposed centromere-proximal inverted repeat (IR) in single-stranded DNA mediates intra-strand foldback and annealing. If the IR is near the DSB, foldback annealing forms a short heterologous flap that is removed by the proof-reading activity of Pol δ. If the IR is further away from the DSB, a longer flap is formed that may be cleaved by Rad1-Rad10. Following flap removal, Pol δ synthesizes from the 3′-end of the foldback, catching up with the resected end. After ligation, a hairpin-capped chromosome is formed with a terminal deletion. During the next cell cycle, the DNA replication machinery loops around the hairpin end and synthesizes back, generating a dicentric isochromosome. If the two centromeres of the isochromosome attached to opposite spindle pole bodies, they are pulled to each nascent daughter cell during mitosis and the chromosome is broken during cytokinesis. The broken chromosome is then stabilized either through the acquisition of a telomere or through centromere loss. The former occurs mainly through break-induced replication initiated by Rad51-mediated strand-invasion from a repeat sequence (such as a delta element) near the broken end into a homologous sequence in another chromosome. Alternatively, the broken chromosome can undergo strand annealing with another chromosome through repeat sequences in a Rad51-independent fashion.

The requirement for the IR in the formation of inverted duplications strongly suggests that a foldback at the repeats is an initiating event. Inter-chromosomal annealing between repeats present on two broken sister-chromatids is expected to be much less efficient than intra-molecular annealing between nearby sequences. Further support for the foldback model comes from the observation that increasing the distance between the DSB and the IR reduces the survival frequency of *sae2*Δ cells. We imagine that the longer heterologous flap formed following annealing between the repeats results in less stable base-pairing.

A consequence of foldback annealing is the presence of a heterologous flap that needs to be cleaved before fill-in synthesis if the break is at a distance from the IR. Rad1-Rad10 is required for cleavage of heterologous flaps that are longer than 20 nt, while the proof-reading activity of Pol δ acts redundantly with Rad1-Rad10 to remove shorter flaps ^53^. Surprisingly, we found that Rad1 is not required for inverted duplications when the DSB is 20 bp away from the IR, whereas *sae2*Δ cells with a defect in the proof-reading activity of Pol δ exhibited a significant defect in the formation of inverted duplications. These data suggest that Pol δ proof-reading plays a non-redundant role in cleaving the short heterologous flaps formed. Although we did not observe a significant decrease in survival of *sae2*Δ *rad1*Δ cells to a DSB induced 51 or 163 bp away from the target IR, the fraction of inverted duplications that initiated from the target IR was lower than in *sae2*Δ cells, consistent with a role for Rad1-Rad10 in removal of long heterologous flaps.

The Rad1-Rad10 complex was also found to be dispensable for spontaneous inverted duplications in *sae2*Δ cells ^7^. However, there was a change in the IRs used for inverted duplications. An IR hotspot with a 3-nt loop and a perfect 15-bp stem was used less frequently, and there was a relative increase in the usage of inverted repeats with imperfect stem loops. We sequenced eight inverted duplication clones from the *sae2*Δ *rad1*Δ double mutants and all were centered on the target IR. The only exceptions were two clones from the *sae2*Δ *rad1*Δ *pol3-01* triple mutant which had used an inverted repeat closer to the DSB, predicted to reduce the size of the heterologous flap after foldback annealing.

The dicentric chromosome predicted to form by replication of the hairpin-capped intermediate would need to be stabilized by loss of one centromere or by acquisition of a telomere following chromosome breakage ^24, 26^. The recovery of two GCRs from *sae2*Δ derivatives with isochromosomes and deletion of one centromere provides strong support for this model. A model for inverted duplication formation without hairpin-capped and dicentric chromosome intermediates has been proposed ^5, 7^. In this model, fill-in synthesis following foldback intra-strand annealing undergoes a template switch at a repeat sequence with a homologous sequence at a different locus, initiating BIR synthesis to the end of the donor chromosome. The requirement for Pol32 shown here would also be consistent with this mechanism and we cannot rule out that it is responsible for some inverted duplications.

There are multiple mechanisms that can lead to telomere acquisition, including *de novo* telomere addition or translocation that leads to telomere capture. Translocations have been shown to mainly occur by HR using repeat elements ^5–7, 25^. In agreement with previous studies ^5–7^, the secondary rearrangements we observed involved repeat sequences. We detected rearrangements mainly using delta elements and *PAU* repeats. Translocations involving *PAU2* have also been detected in spontaneous inverted duplications ^7, 32, 61^. The *PAU2* sequence is located ∼30 kb centromeric to the DSB and ∼90 kb from the centromere. Previous work showed breakage of dicentric chromosomes occurs within a 25-30 kb window of the centromere ^22, 23^. *YELCDELTA4* is located ∼14 kb away from the centromere, which falls in the middle range of the breakage window. Consistently, 16/37 inverted duplications associated with secondary recombination events used *YELCDELTA4*. The next δ element, *YELCDELTA3*, is 50 kb from the centromere but is much shorter than other LTRs (174 bp vs ∼300-360 bp) and was used in 12 inverted duplications. *PAU2,* which is located ∼30 kb centromeric to the DSB and ∼90 kb from the centromere, was utilized in the remaining 9 inverted duplications. Since more than half of the inverted duplications sequenced used *YELCDELTA3* or *PAU2*, the majority of breakage events likely occur further telomeric from *YELCDELTA4.* The bias for use of delta elements is likely due to the high copy number of these sequences providing many potential templates for repair.

Since breakage of the dicentric chromosome is unlikely to occur within the repeats used for secondary events, Rad51-dependent strand invasion would require removal of heterologous flaps. Given the role of Rad1 in heterologous flap cleavage during HR and SSA ^51^, the high survival frequency of *sae2*Δ *rad1*Δ and use of a repeat sequence for the secondary events is surprising. However, two of the inverted duplications recovered from the *sae2*Δ *rad1*Δ mutant had complex rearrangements and one was associated with telomere addition, suggesting that stabilization of a broken dicentric intermediate by HR might be more problematic in cells lacking Rad1-Rad10 nuclease.

The long homology involved in the secondary rearrangements and non-reciprocal nature of the rearrangements point to a BIR mechanism or a crossover in which only one product is recovered in a daughter cell. Further support for either mechanism is the significant reduction in survival frequency of *rad51*Δ *sae2*Δ compared to *sae2*Δ cells. This indicates that the secondary rearrangements involve a strand invasion step. The survivors with inverted duplications detected in *sae2*Δ *rad51*Δ cells are associated with homology-mediated secondary events, potentially arising by SSA involving another broken chromosome (Figure 7). The increase in spontaneous lesions in *rad51*Δ may allow for the detection of rearrangements using these other mechanisms by increasing the likelihood of the presence of a second broken chromosome.

The role of BIR in the secondary rearrangements in spontaneous inverted duplications has been questioned based on the lack of effect of *pol32*Δ on the rate of these events ^7^. While Pol32 has in fact been shown to be required for BIR ^57^, deletion of *POL32* may have pleiotropic effects on GCR that confound interpretation of its role in inverted duplications. Perhaps *POL32* deletion increases the frequency of the initiating substrates, which in turns has more of an effect on the rate of inverted duplication than it does on altered repair pathway. Spontaneous GCRs with non-reciprocal translocations have been observed in the *pol32*Δ mutants before ^4^. Indeed, *pol32*Δ cells have overall higher GCR rates compared to wild-type cells ^4^. Therefore, in the GCRs discussed above, while the translocations could have occurred by BIR, another mechanism for the formation cannot be ruled out. Interestingly, we recovered fewer inverted duplications in *sae2*Δ *pol32*Δ cells than in *sae2*Δ *rad51*Δ cells, suggesting an additional role for Pol32 in formation of inverted duplications. Since a previous study reported fewer DSB-induced hairpin-capped chromosomes in RPA-depleted *pol32*Δ cells ^56^, this function is likely to be at the fill-in synthesis step after foldback annealing and heterologous flap removal.

Foldback inversions have been detected in a variety of human cancers, including pancreatic, ovarian, breast, and squamous cell carcinomas (SCC), and are associated with poor prognosis ^62–68^. The mechanism by which these foldback inversions form is unknown and could involve the foldback priming mechanism described here, particularly in tumors with mutations in *MRE11, RAD50* or *NBS1*. However, we note that mammalian cells have more active end joining mechanisms than yeast ^69^; therefore, fusion of broken sister chromatids is likely to contribute significantly to the generation of foldback inversions in human cells.

## MATERIALS AND METHODS

### Strains, media, and growth conditions

Yeast strains used in this study are listed in Table S2. Yeast strains were made either by crossing or transformation. Transformation was performed using the LiAc/ssDNA carrier DNA/PEG method ^70^. Cells were grown in either yeast rich media (YPD) or synthetic complete (SC) media with amino acid dropouts as described previously ^71^. All cells were propagated at 30°C. β-estradiol containing plates were made by spreading β-estradiol on YPD plates to 5 µM before use. pAA3, pAA13, pAA16, pAA18, pAA19, pAA20, pAA21 and pAA22 were integrated into the yeast genome by digesting the plasmids with AscI and transforming cells with the linearized product. Integration was confirmed by multiplex PCR as described previously ^72^.

Complete deletion of the *RAD1, RAD51,* and *POL32* ORFs was achieved by one-step gene disruption. For each, a PCR fragment was made by amplifying either HphMX (*rad1*Δ) or NatMX (*rad51*Δ and *pol32*Δ) from pAG32 (Addgene plasmid # 35122) or pAG25 (Addgene plasmid # 35121), respectively ^73^. Forward (TCGACGGATCCCCGGGTTAA) and reverse (AATTCGAGCTCGTTTTCGACACT) primers contained 60 bp of sequences immediately upstream and downstream of the ORFs, respectively. Yeast cells were then transformed with the fragment, plated on YPD and then replica plated on the appropriate selection media after one day of growth. Gene deletion was confirmed by multiplex PCR.

The inverted repeat was mutated by CRISPR/Cas9 gene editing as described previously^74^. The gRNA IR_gRNAmut2 was cloned into pCeASY and used for editing (Table S3). The repair template included the desired mutations along with 500 bp of homologies telomeric (amplified using IR_500_upstream_F and mutation-specific reverse primer) and centromeric to the target site (amplified using IR_500_downstream_R and a mutation-specific forward primer) and was assembled by overlapping PCR using Phusion polymerase. The targeted modifications were confirmed by sequencing.

The *pol3-01* allele was made by CRISPR/Cas9 gene editing as described above. The repair template containing the *pol3-01* mutation ^54^ along with ∼75 bp of upstream and downstream homology sequences was assembled by PCR using the primers pol3-01_middle, pol3-01_left and pol3-01_right, listed in Table S3. The mutation was confirmed by sequencing of DNA amplified by pol3-01_mut_F and pol3-01_mut_R.

### Plasmids and constructs

The oligonucleotides used for gRNAs and genome editing are listed in Table S3, and plasmids are listed in Table S4. Details of the oligonucleotides used for plasmid modifications and assembly are available on request.

#### Cas9 expression plasmids

pAA1 was made by cloning the *GAL1* promoter into pML107 (Addgene plasmid # 67639)^75^. The *GAL1p* promoter was amplified by PCR from genomic DNA and assembled into NcoI and SpeI digested pML107 using the HiFi DNA Assembly mix (NEB # E5520). The gRNA scaffold sequence was further modified to replace the SwaI and BclI fragment with a BaeI fragment from pML107 by HiFi Assembly.

pAA3 was made by cloning *P_GAL1_-CAS9-NLS-6xGLY-FLAG-T_CYC1_* into pRG203MX ^72^. *P_GAL1_-CAS9-NLS* was amplified from pAA1 and the *CYC1* terminator was amplified from genomic DNA. The two fragments were then assembled into SpeI- and EcoRV-digested pRG203MX.

The lexO expression system is a two-module system composed of the promoter and the transcription factor. The promoter is composed of four tandem repeats of the lexA box fused to the minimal *CYC1* promoter (together referred to as *P_lexO_*) ^36^. The transcription factor is LexA gene fused to an estrogen receptor and a B112 transcriptional activator under the control of *ACT1* promoter (together referred to as *P_ACT1_-LexA-ER-AD* ^36^. The two modules were cloned into a single plasmid to generate pAA12, which was then used for subsequent cloning. pAA16 was made by amplifying *CAS9-NLS-FLAG* from pAA3 and was cloned into ApaI-SacI-digested pAA12 by HiFi assembly. pAA18 was made by HiFi assembly. *CAS9* (amplified from pAA3), ER-LBD (amplified from pRG634) and NLS-FLAG (amplified from pAA3) were assembled with ApaI-SacI-digested pAA12.The orientations of *P_lexO_-CAS9-NLS-FLAG-T_ADH1_* and *P_lexO_-CAS9-NLS-FLAG-ER-T_ADH1_* were reversed relative to *P_ACT1_-LexA-ER-B112-T_CYC1_* by re-cloning the NotI fragment of pAA19 and pAA20 respectively, into pAA12.

#### Integrating gRNA constructs

A version of pCAS (Addgene plasmid # 60847) ^74^ in which the gRNA cloning sequences were replaced with an XbaI and ZraI fragment was kindly provided by R. Gnügge. The sgRNA from this plasmid, including the tRNA promoter, the HDV ribosome, the sgRNA sequence and the sNR52 terminator, was PCR amplified and cloned into ApaI- and XhoI-digested pRG205MX ^72^ whose backbone XbaI and ZraI sites were then destroyed by site-directed mutagenesis. This vector was named pAA9.

The gRNA targeting sequences were cloned into pAA9 by annealing oligos pCeASY-gRNA-S and pCeASY-gRNA-AS (where 20xN represent the sense and anti-sense gRNA target, respectively) and ligating the annealing product to ZraI- and XbaI-digested pAA9.

#### Cas9-mediated gene editing plasmids

pCeASY, a version of pCAS (Addgene plasmid # 60847) ^74^ in which the gRNA cloning sequences were replaced with an XbaI and ZraI fragment was kindly provided by R. Gnügge. gRNA targeting sequences were cloned into this plasmid as described above.

### Survival assays

All survival assays with *lexO-CAS9-ER* were done in strains in which the relevant gRNA was integrated into the genome. Fresh single colonies from each strain were grown overnight in YPD, diluted in the morning and grown to early log phase. Cells were then diluted and plated on YPD ± β-estradiol. The survival frequency was measured by: 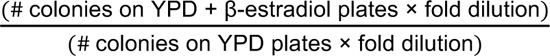

### Screening of survivors for mutagenic repair

PCR and qPCR screens were done using DNA extracted from boiled cells. Colonies were picked from the survival assay plates, spread onto patches on YPD plates and grown overnight. A colony-sized amount of cells was then boiled in 50 µl 0.2% SDS at 95°C for 5 min. All screening PCR reactions were performed using DreamTaq (ThermoFisher # EP0711) in the presence of 1% Triton X-100 and using 1 µl of boiled cell lysates in a 20 µl reaction. Control primers MEC1-F2 and MEC1-R2 were used in the same PCR tubes for all reactions (Table S2). The PCR screen to detect NHEJ products was done using primers P3 and P4. For screening for retention of the terminal part of the left arm of chromosome V, the same PCR reaction was used but with primers P1 and P2 instead of the P3/P4 primers. WT clones for which there was no P3/P4 but which gave P1/P2 product were further screened using primer set P7 and P8 to detect homeologous gene conversion events that lead to loss of P4 priming sequences. Primer sets P11/P12 and P13/P14 were used to screen survivors from gRNA-48 and gRNA-160 expression for use of the target IR. Clones that were negative for P11/P12 and positive for P13/14 were scored at target IR inverted duplications, clones that were positive for both diagnostic PCRs were scored as using an IR telomeric to the target IR, and clones that were negative for both PCRs were scored as using an IR centromeric to the target IR. Differences in IR usage between *sae2*Δ and *sae2*Δ *rad1*Δ were determined by Fisher’s exact test. We identified a sub-population of fast-growing survivors from WT and *sae2*Δ *pol32*Δ cells that had no alteration at the gRNA cut site. These were shown by a PCR assay to have inverted the DNA segment between the ER domains in the expression construct resulting in failure to express Cas9.

To screen for inverted duplications, DNA was prepared from cells that were serially passaged for a total of two times (from survival plate to “patch 1,” then from “patch 1” to “patch 2”). Boiled DNA (as above) was diluted 10-fold in 2% Tween 20, of which 4.4 µl was used in a 10 µl SYBR Green qPCR reaction. Primer set ChrV-60K-F and ChrV-60K-R (P5 and P6) were used to probe for the duplication on Chr V, and ADH1-fwd and AHD1-rev were used for a reference amplicon. Each reaction was run in triplicates. During each qPCR run, two parental controls were run to normalize the sample Cq values to. The copy number was measured by: 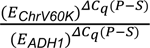 were *E_ChrV60K_* and *E_ADH1_*are the primer efficiencies for the *ChrV-60K* and *ADH1* amplicons, respectively, and *ΔC_q_(P − S)* is the difference between the quantification cycles of the parental and surviving clones.

### Pulse-field gel electrophoresis (PFGE)

DNA for PFGE was extracted in low-melting point agarose from cells grown to saturation as described previously ^76^. The chromosomes were separated in a BioRad CHEF-DR II. The gel was then stained using SYBR Gold (Invitrogen # S11494) for one hour. The chromosomes from the gel were transferred to nylon membranes (Hybond N+) and hybridized to radioactively labeled *PCM1* to identify Chr V fragments.

### Southern blotting to detect inverted duplications

3-5 µg of DNA isolated from surviving clones was digested for eight hours with 15 units of the indicated restriction enzymes, separated on agarose gels and transferred to nylon membranes (Hybond N+). The DNA was then hybridized with breakpoint-specific radiolabeled probe to detect inverted duplications.

### Whole genome sequencing

Genomic DNA for library preparation was isolated using YeaStar Genomic DNA Kit (Zymo Research # D2002). DNA libraries were prepared using the Illumina DNA Library Prep (Illumina # 20018704). For each sample 100 ng of genomic DNA was used following the manufacturer’s instructions with the following exceptions: we used half the recommended volume of each reagent, and we modified the size selection scheme to achieve larger fragments as follows: amplified tagmented DNA was diluted 1:4.44 in water (22.5uL DNA in 77.5uL water). During the first size selection step, 44.7uL SPB was added to diluted DNA. After mixing and incubation, 142uL of the supernatant was used during the second size selection step with 10uL SPB. Each library was resuspended in 30 µl buffer and the libraries were then pooled in equimolar amounts. After pooling, a final 0.4x SPB size selection (40uL SPB added to 50uL pool + 50uL water) was performed in order to eliminate small fragments prior to submitting the pool for sequencing. Pooled libraries were diluted and denatured according to the manufacturer’s recommendations. Paired-end read sequencing was done in a NextSeq 500/550 platform using either the 150-cycle Medium Output Kit (Illumina # 20024909), 75 cycles per read, or the 75 cycle High Output Kit (Illumina # 20024906), 37 cycles per read.

All mapping was done to a W303 reference genome ^77^. For all analysis, all reads where quality filtered with fastp using default settings before mapping ^78^. For copy number variation analysis (CNV), the paired-end reads were mapped using Bowtie2 ^79^. Samtools ^80^ was used to remove PCR duplicates and to extract sequencing depths for each position along the genome for each sample. CNV was calculated relative to a parental clone that was sequenced with the GCR samples. Briefly, for each the parental and the GCR clone, the number of reads mapping to each position was normalized to the total number of reads. Next, the chromosome positions were binned (either in 5 kb or 1 kb windows, depending on the analyses shown in the figures). Finally, the relative copy number was calculated for each bin by taking the ratio of the normalized number of reads for each bin from the GCR clone to that of the parental clone. A constant of 0.0625 was added to each computed CNV value to allow for *log_2_* transformation. For significant value calculation, first, the same CNV analysis was done by randomly splitting the parental reads into two and finding the CNV between the split samples. Then, the CNV values for the split parental samples were used as the null distribution, which was then used to determine the *p* value of the CNV for each bin of the GCR sample. Finally, multiple testing correction was done using the false discovery rate method. The threshold for significance was set at 0.001.

Structural variation (SV) was detected using Comice, part of the Pyrus suite ^5^. The reads for each sample were aligned separately using Bowtie2 and then processed using Comice. The sequence at the center of the SVs was obtained using two strategies: the first was from the outcome of Comice, the second was form *de novo* assembly of the discordant reads near the boundary of the SV variants. For the second strategy, SV boundaries were determined based on the change CNV data. Next, discordant reads pairs for which one mate pair maps to within a few kb of the SV boundary were extracted from the mapped BAM files using samtools. The extracted reads were then *de novo* assembled using Unicycler ^81^. WGS data is publicly available at the following website: http://www.ncbi.nlm.nih.gov/bioproject/900608.

## ACKNOWLEDGMENTS

We thank C. Putnam for advice on WGS analysis and members of the Symington lab for review of the manuscript. This work was supported by grants from the National Institutes of Health (P01 CA174653 and R35 GM126997 to L.S.S., and in part through the NIH/NCI Cancer Center Support Grant P30CA013696 [CCSG DNA Sequencing Core]).

## AUTHOR CONTRIBUTIONS

A.A. performed most of the experiments shown in Figures 1-6 and S1-5; M.R.N. contributed to the experiments in Figures 1, 4-6 and S5. A.A., M.R.N. and L.S.S. contributed to the study design, data analysis and manuscript preparation.

## SUPPLEMENTARY FIGURE LEGENDS

**Figure S1.**
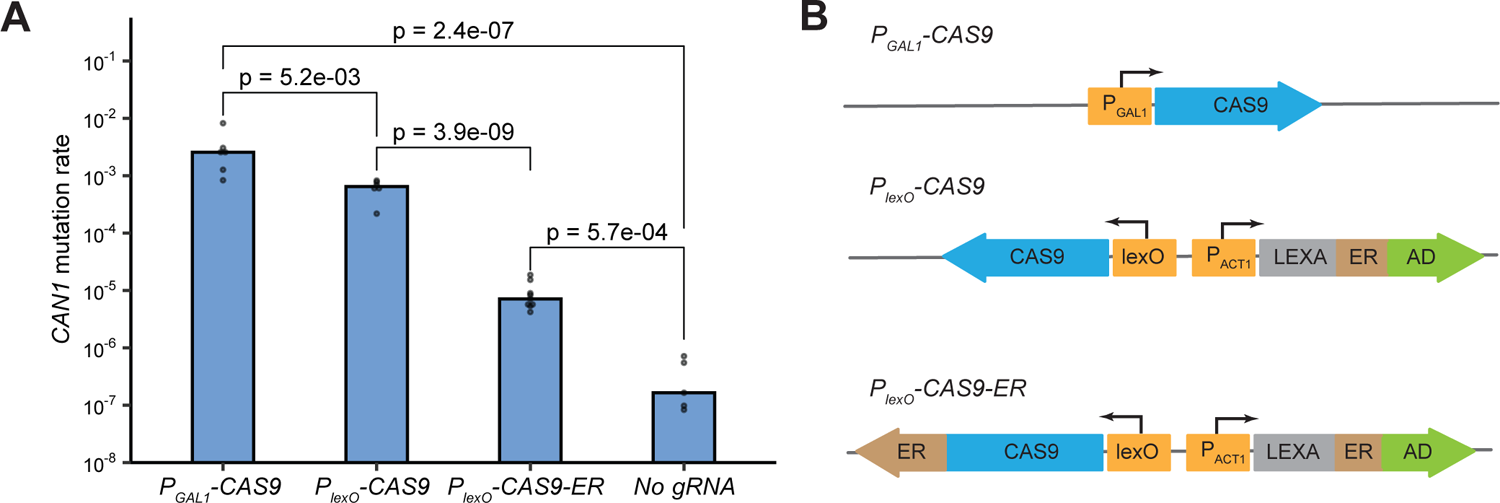
Leaky expression of Cas9. A. Spontaneous mutation rate of the *CAN1* locus in WT cells carrying the indicated the *CAS9* constructs with a gRNA targeting the *CAN1* locus or without gRNA. Cas9 expression was not induced during the experiment. B. Constructs used to express Cas9.

**Figure S2.**
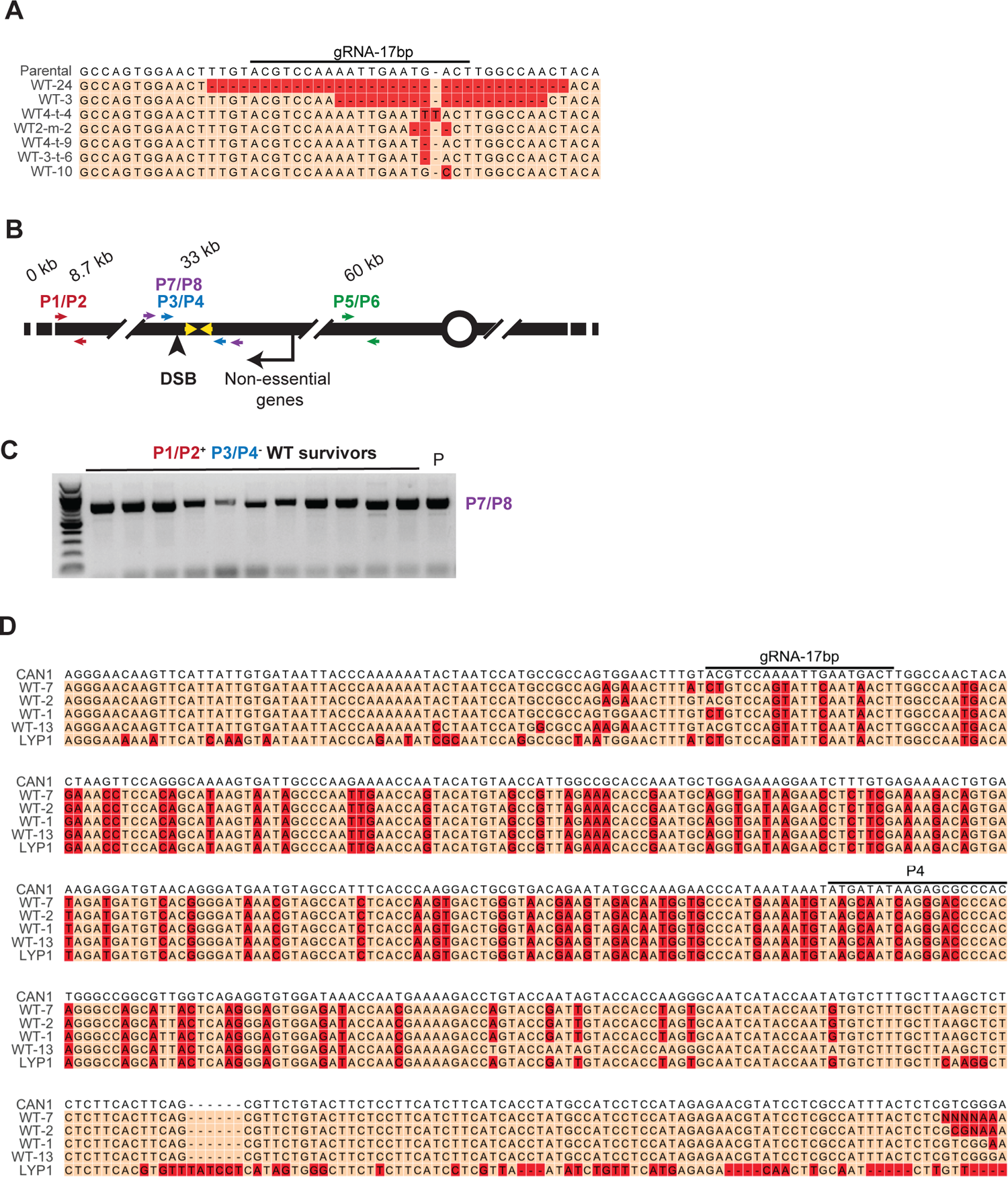
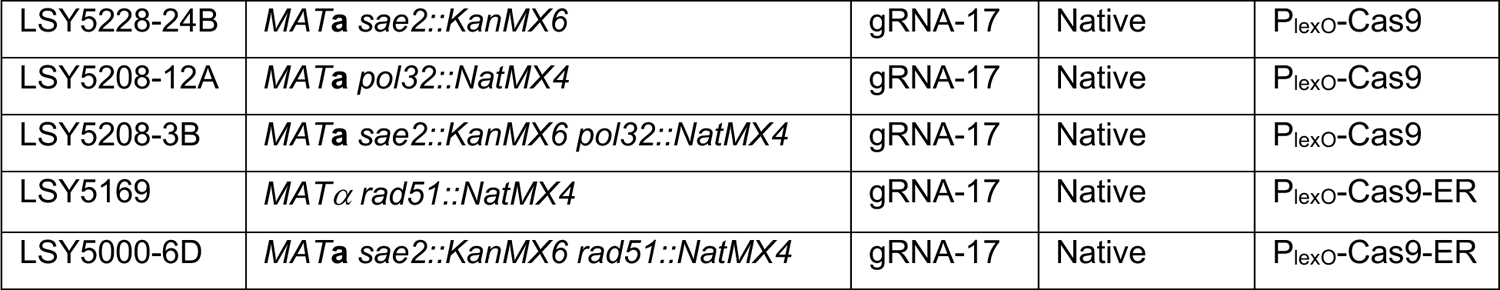
Molecular analysis of survivor types from WT cells. A. The P3/P4 primer PCR products from select WT clones were sequenced and show evidence of indels. B. Schematic of the left arm of Chr V showing the location of the DSB and inverted repeats (yellow arrows). Primer pairs to detect retention of terminal sequence, NHEJ events and *LYP1* conversions are shown in red, blue and purple, respectively. C. Clones that had P1/2 but not P3/4 PCR bands were amplified using the P7/P8 primer pair to detect the presence of sequences further away from P3/P4. D. Sanger sequence of P7/P8 primer PCR products shows evidence of *LYP1* conversions.

**Figure S3.**
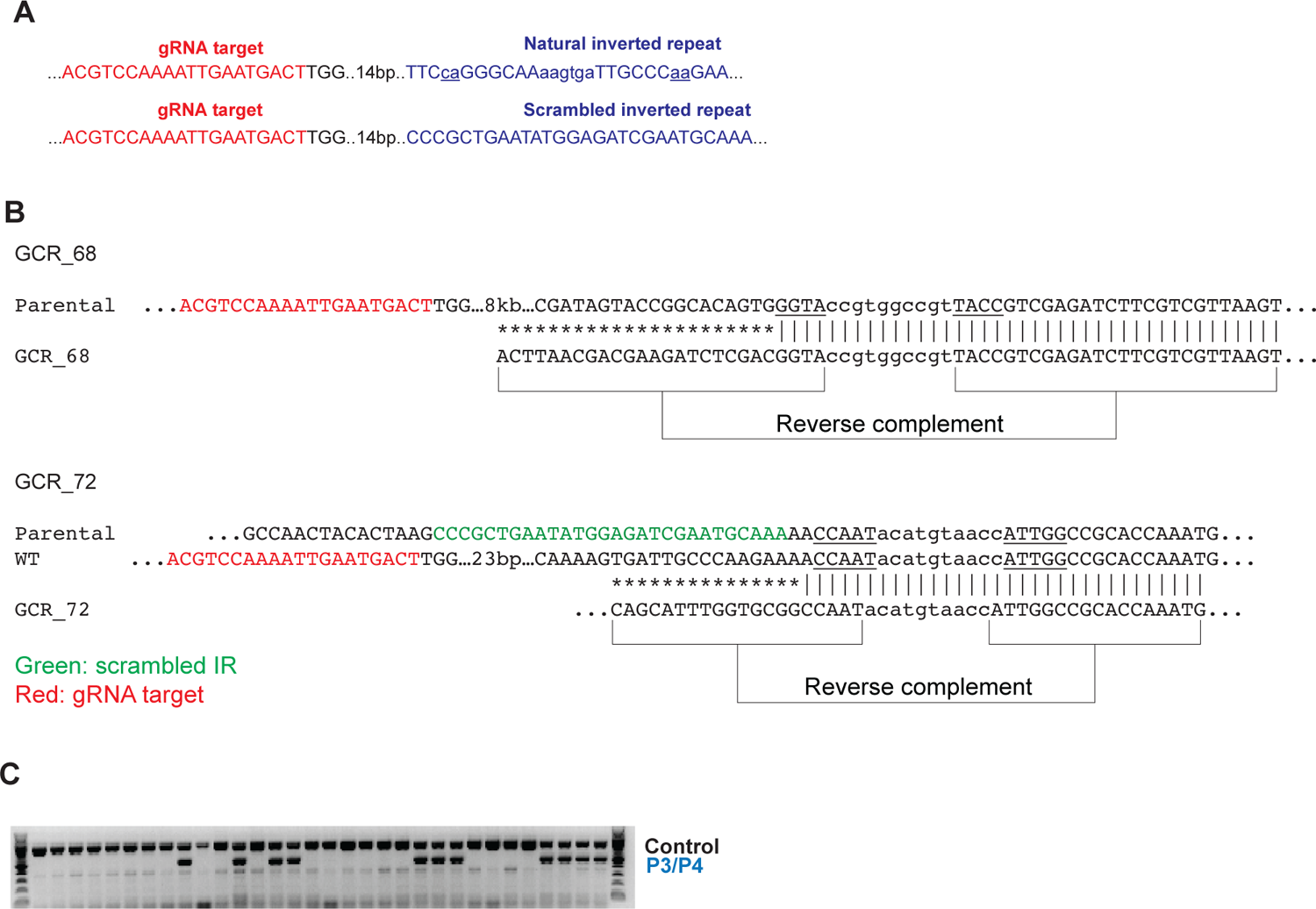
Analysis of *sae2*Δ survivors from the scrambled IR strain. A. Sequence of the original IR and scrambled IR located 17 bp from the gRNA target sequence. B. The sequence at the center of the inverted duplications of two *sae2*Δ clones with the scrambled IR. Mismatches in the inverted repeat are shown in lower case. In red is the gRNA target sequence. For GCR_72, the position of the scrambled IR within the unrearranged sequence is shown for reference. C. 32 independent clones from *sae2* cells with scrambled inverted repeat were analyzed using P3/4 primers to detect NHEJ events.

**Figure S4.**
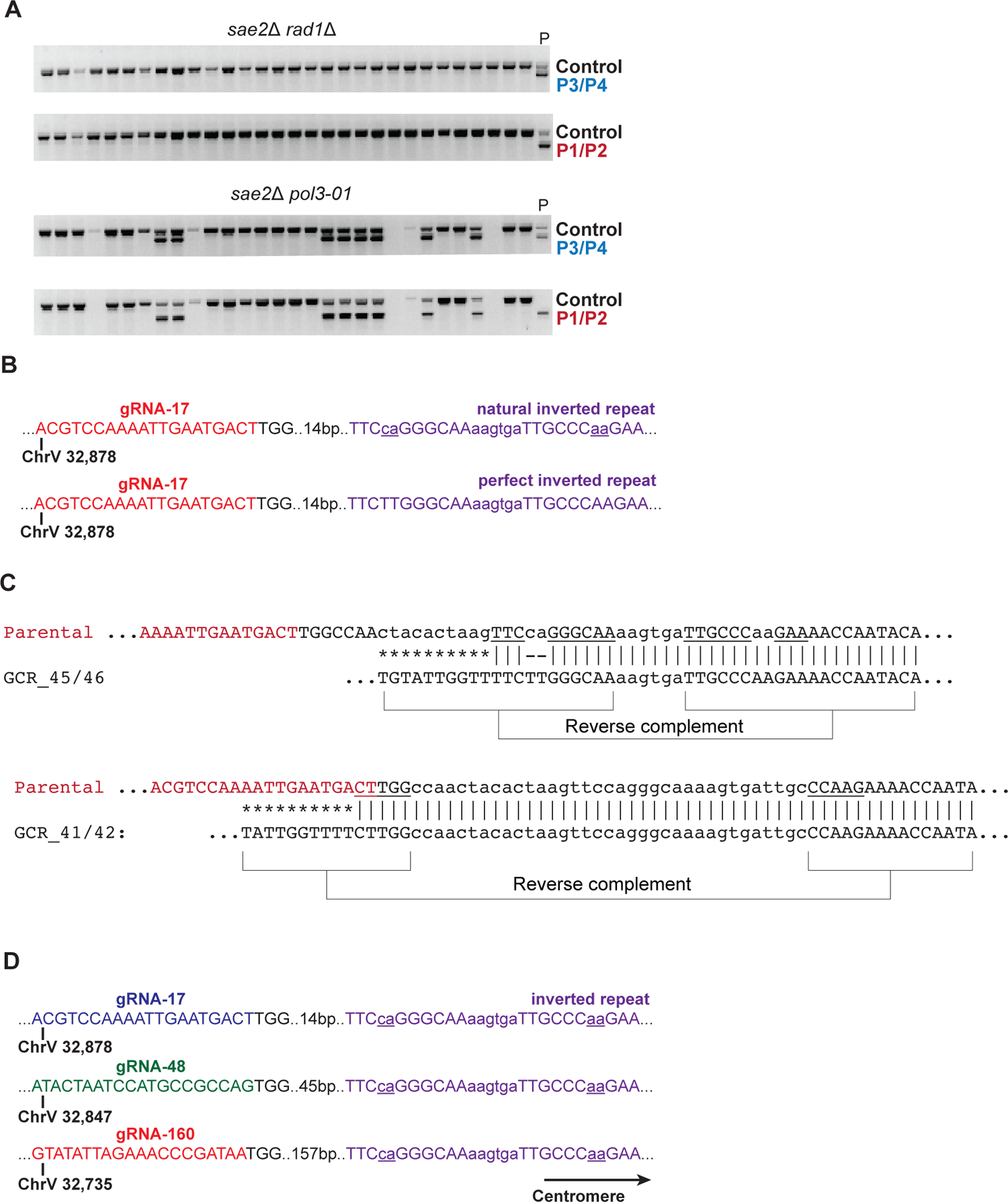
Analysis of *sae2*Δ *rad1*Δ and *sae2*Δ *pol3-01* clones. A. 30 independent clones from *sae2*Δ *rad1*Δ and *sae2*Δ *pol3-01* cells were analyzed using P3/4 and P1/2 primers to detect NHEJ events or loss of Chr V terminal sequence. “P” denotes the parental strain. B. The sequence of the inverted repeat (purple font) mutated to correct the mismatches (underlined, lower case). C. GCR_45 and GCR_46 have the target IR (shown underlined in the unrearranged sequence above) as the center of the inverted duplications, while GCR_41 and GCR_42 used an IR directly adjacent the cut site. Note: the latter IR is the same used in the inverted duplications in WT clones (Figure 2F). D. The position of the gRNA-17, gRNA-48 and gRNA-160 relative to the inverted repeat.

**Figure S5.**
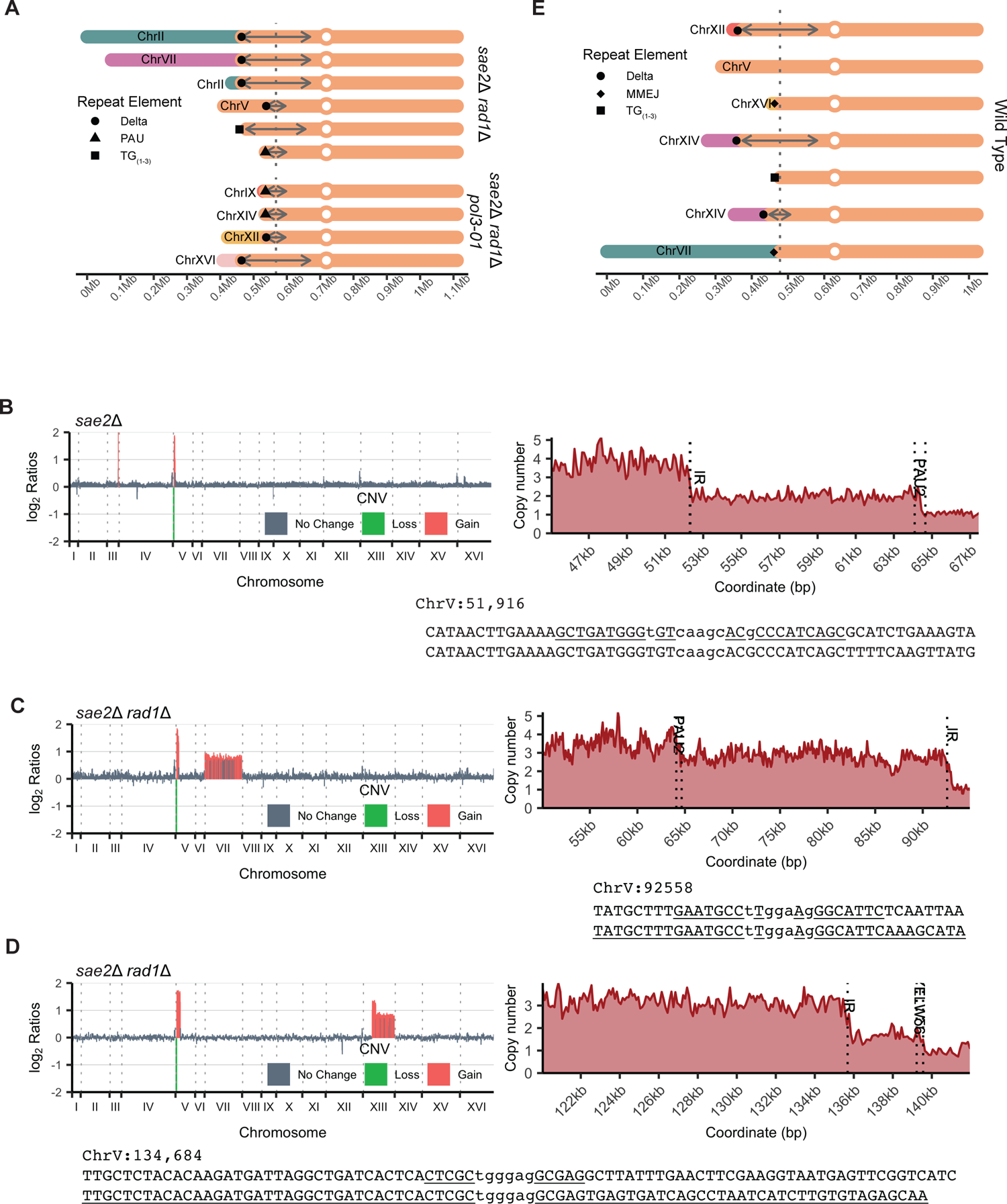
NGS analysis of inverted duplication clones. A. Derivative Chr V in *sae2*Δ *rad1*Δ and *sae2*Δ *rad1*Δ *pol3-01* clones. B-D. Clones that exhibit quadruplications centromeric to the target IR in a *sae2*Δ (B) and in two *sae2*Δ *rad1*Δ clones (C and D). Left: log_2_ ratios of genome-wide copy number relative to parental reads. Right: relative copy number of junctions between the higher order duplication order copy number and duplication sequence, and between the latter and non-duplication sequence. In each case, a naturally occurring IR occurs at the junction of the drop in copy number. Below each panel is the IR sequence highlighted in the right panels and occur at the center of an inverted duplication. E. Derivative Chr V in WT clones.

---

**Table S1.**
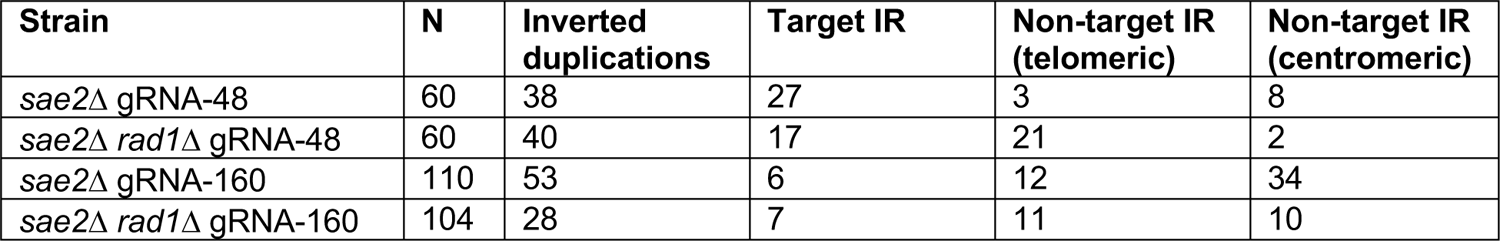
Analysis of survivors using gRNA-48 and gRNA-160

| **Strain** | **N** | **Inverted duplications** | **Target IR** | **Non-target IR (telomeric)** | **Non-target IR (centromeric)** |
| --- | --- | --- | --- | --- | --- |
| *sae2Δ* gRNA-48 | 60 | 38 | 27 | 3 | 8 |
| *sae2Δ rad1Δ* gRNA-48 | 60 | 40 | 17 | 21 | 2 |
| *sae2Δ* gRNA-160 | 110 | 53 | 6 | 12 | 34 |
| *sae2Δ rad1Δ* gRNA-160 | 104 | 28 | 7 | 11 | 10 |

**Table S2.** Yeast Strains

| Strain | Relevant Genotype<sup>1</sup> | gRNA | IR | Cas9 |
| --- | --- | --- | --- | --- |
| W1588-4C | *MATa* | n/a | Native | n/a |
|  |  | gRNA-17 | Native | P<sub>GAL</sub>-Cas9 |
| LSY4708 | *MATa* | n/a | Native | P<sub>lexO</sub>-Cas9-ER |
| LSY4713 | *MATa* | gRNA-17 | Native | P<sub>lexO</sub>-Cas9-ER |
| LSY4714 | *MATα* | gRNA-17 | Native | P<sub>lexO</sub>-Cas9-ER |
| LSY4751-11C | *MATα sae2::KanMX6* | gRNA-17 | Native | P<sub>lexO</sub>-Cas9-ER |
| LSY4752-4C | *MATα mre11::HIS3MX6* | gRNA-17 | Native | P<sub>lexO</sub>-Cas9-ER |
| LSY4763-7D | *MATa mre11-H125N* | gRNA-17 | Native | P<sub>lexO</sub>-Cas9-ER |
| LSY4903 | *MATa* | gRNA-17 | Scrambled | P<sub>lexO</sub>-Cas9-ER |
| LSY4913-2D | *MATα sae2::KanMX6* | gRNA-17 | Scrambled | P<sub>lexO</sub>-Cas9-ER |
| LSY4616 | *MATa rad1::HphMX6* | n/a | Native | n/a |
| LSY4762-6A | *MATα rad1::HphMX6* | gRNA-17 | Native | P<sub>lexO</sub>-Cas9-ER |
| LSY4762-45D | *MATα sae2::KanMX6 rad1::HphMX4* | gRNA-17 | Native | P<sub>lexO</sub>-Cas9-ER |
| LSY5068-8D | *MATa sae2::KanMX6 pol3-01* | gRNA-17 | Native | P<sub>lexO</sub>-Cas9-ER |
| LSY5068-14D | *MATα sae2::KanMX6 pol3-01*<br>*rad1::HphMX4* | gRNA-17 | Native | P<sub>lexO</sub>-Cas9-ER |
| LSY5052 | *MATa* | gRNA-48 | Native | P<sub>lexO</sub>-Cas9-ER |
| LSY5054 | *MATα sae2::KanMX6* | gRNA-48 | Native | P<sub>lexO</sub>-Cas9-ER |
| LSY5056 | *MATa rad1::HphMX4* | gRNA-48 | Native | P<sub>lexO</sub>-Cas9-ER |
| LSY5193 | *MATα sae2::KanMX6 rad1::HphMX4* | gRNA-48 | Native | P<sub>lexO</sub>-Cas9-ER |
| LSY5053 | *MATa* | gRNA-160 | Native | P<sub>lexO</sub>-Cas9-ER |
| LSY5055 | *MATα sae2::KanMX6* | gRNA-160 | Native | P<sub>lexO</sub>-Cas9-ER |
| LSY5057 | *MATa rad1::HphMX4* | gRNA-160 | Native | P<sub>lexO</sub>-Cas9-ER |
| LSY5059 | *MATa sae2::KanMX6 rad1::HphMX4* | gRNA-160 | Native | P<sub>lexO</sub>-Cas9-ER |
| LSY4904 |  | gRNA-17 | Perfect | P<sub>lexO</sub>-Cas9-ER |
| LSY5171-23B | *MATα sae2::KanMX6* | gRNA-17 | Perfect | P<sub>lexO</sub>-Cas9-ER |
| LSY5171-44C | *MATα sae2::KanMX6 rad1::HphMX4* | gRNA-17 | Perfect | P<sub>lexO</sub>-Cas9-ER |
| LSY5171-46A | *MATa sae2::KanMX6 pol3-01* | gRNA-17 | Perfect | P<sub>lexO</sub>-Cas9-ER |
| LSY5235-1 | *MATα sae2::KanMX6 pol3-01*<br>*rad1::HphMX4* | gRNA-17 | Perfect | P<sub>lexO</sub>-Cas9-ER |
| LSY4880 | *MATa pol32::NatMX* | n/a | Native | P<sub>lexO</sub>-Cas9-ER |
|  | *MATa sae2::KanMX6 pol32::NatMX4* | gRNA-17 | Native | P<sub>lexO</sub>-Cas9-ER |
| LSY5228-1A | *MATa* | gRNA-17 | Native | P<sub>lexO</sub>-Cas9 |
| LSY5228-24B | *MATa sae2::KanMX6* | gRNA-17 | Native | P<sub>lexO</sub>-Cas9 |
| LSY5208-12A | *MATa pol32::NatMX4* | gRNA-17 | Native | P<sub>lexO</sub>-Cas9 |
| LSY5208-3B | *MATa sae2::KanMX6 pol32::NatMX4* | gRNA-17 | Native | P<sub>lexO</sub>-Cas9 |
| LSY5169 | *MATα rad51::NatMX4* | gRNA-17 | Native | P<sub>lexO</sub>-Cas9-ER |
| LSY5000-6D | *MATa sae2::KanMX6 rad51::NatMX4* | gRNA-17 | Native | P<sub>lexO</sub>-Cas9-ER |

**Table S3.**
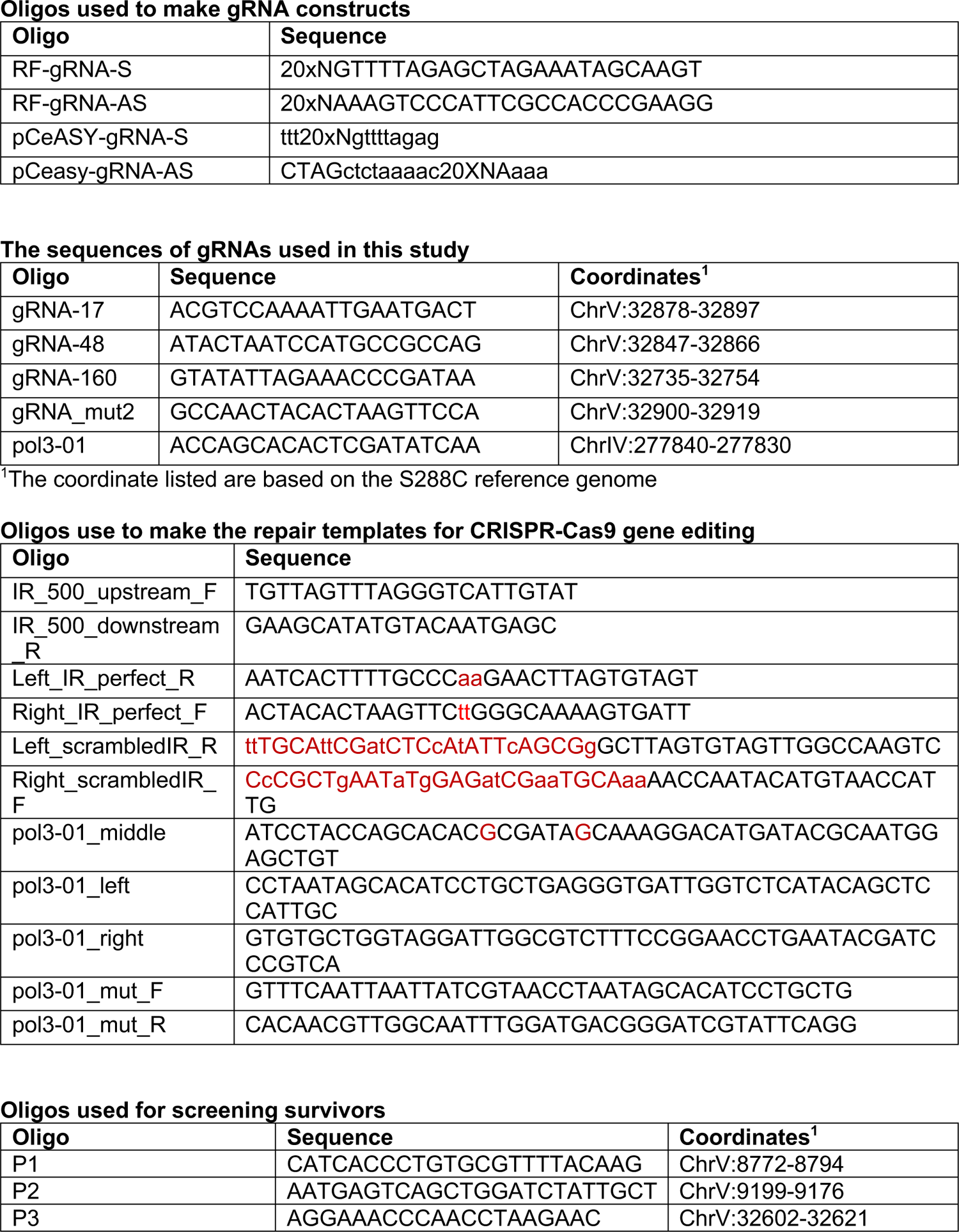
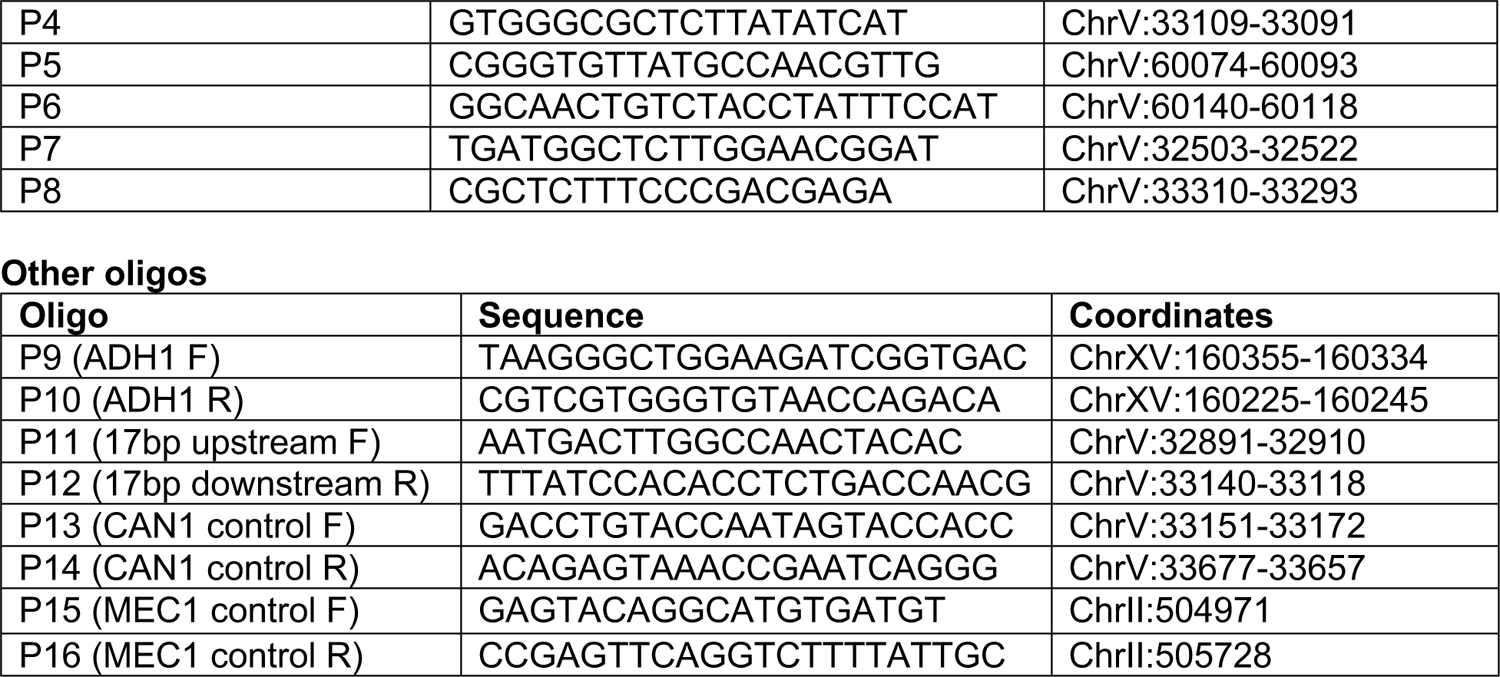
Oligonucleotides

**Table S4.**
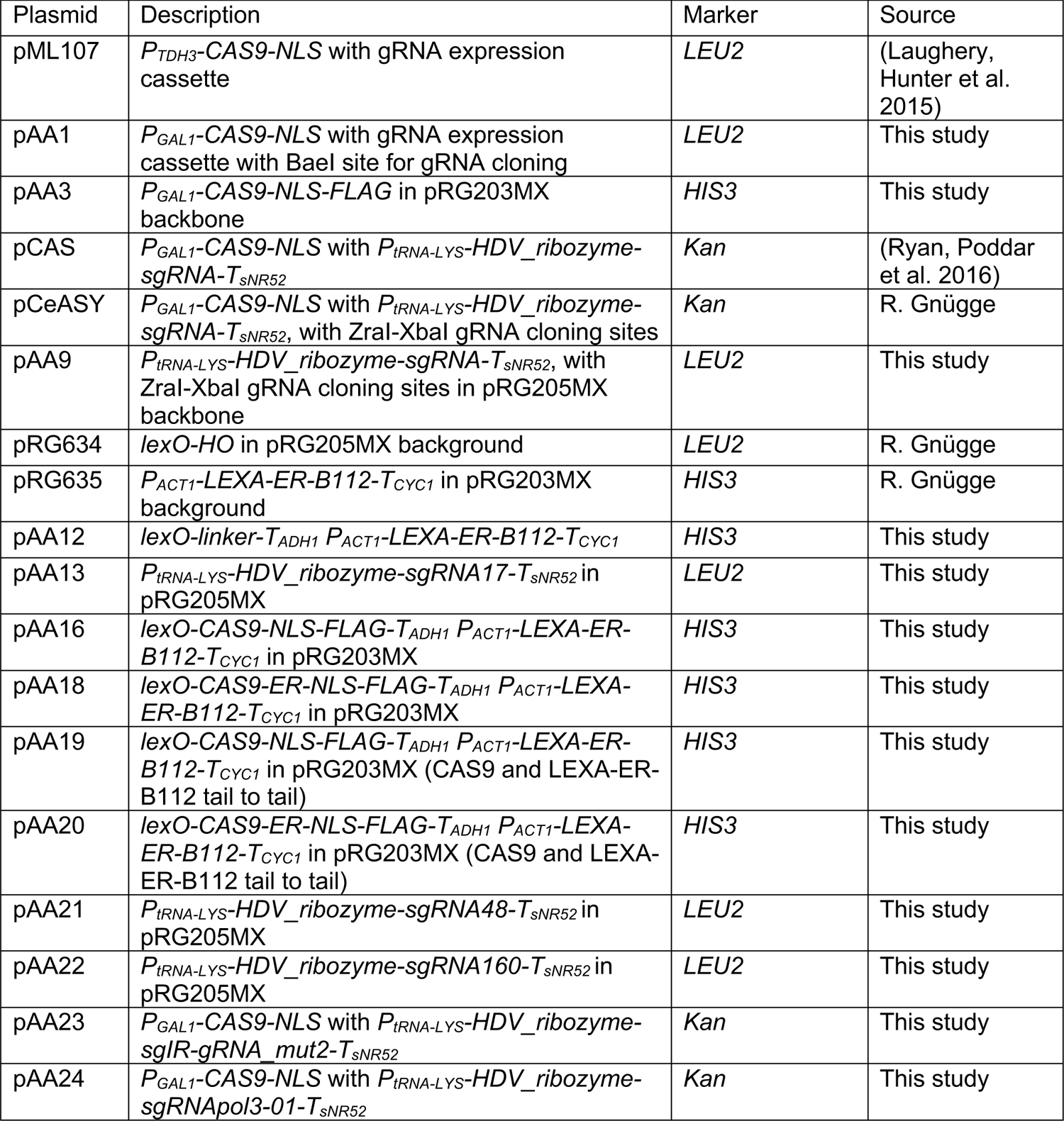
Plasmids used in this study

| Plasmid | Description | Marker | Source |
| --- | --- | --- | --- |
| pML107 | *P<sub>TDH3</sub>*-CAS9-NLS with gRNA expression cassette | *LEU2* | (Laughery, Hunter et al. 2015) |
| pAA1 | *P<sub>GAL1</sub>*-CAS9-NLS with gRNA expression cassette with Bael site for gRNA cloning | *LEU2* | This study |
| pAA3 | *P<sub>GAL1</sub>*-CAS9-NLS-FLAG in pRG203MX backbone | *HIS3* | This study |
| pCAS | *P<sub>GAL1</sub>*-CAS9-NLS with *P<sub>tRNA-LYS</sub>*-HDV_ribozyme-sgRNA-*T<sub>SNR52</sub>* | *Kan* | (Ryan, Poddar et al. 2016) |
| pCeASY | *P<sub>GAL1</sub>*-CAS9-NLS with *P<sub>tRNA-LYS</sub>*-HDV_ribozyme-sgRNA-*T<sub>SNR52</sub>*, with Zral-Xbal gRNA cloning sites | *Kan* | R. Gnügge |
| pAA9 | *P<sub>tRNA-LYS</sub>*-HDV_ribozyme-sgRNA-*T<sub>SNR52</sub>*, with Zral-Xbal gRNA cloning sites in pRG205MX backbone | *LEU2* | This study |
| pRG634 | *lexO-HO* in pRG205MX background | *LEU2* | R. Gnügge |
| pRG635 | *P<sub>ACT1</sub>*-LEXA-ER-B112-*T<sub>CYC1</sub>* in pRG203MX background | *HIS3* | R. Gnügge |
| pAA12 | *lexO-linker-T<sub>ADH1</sub>* *P<sub>ACT1</sub>*-LEXA-ER-B112-*T<sub>CYC1</sub>* | *HIS3* | This study |
| pAA13 | *P<sub>tRNA-LYS</sub>*-HDV_ribozyme-sgRNA17-*T<sub>SNR52</sub>* in pRG205MX | *LEU2* | This study |
| pAA16 | *lexO*-CAS9-NLS-FLAG-*T<sub>ADH1</sub>* *P<sub>ACT1</sub>*-LEXA-ER-B112-*T<sub>CYC1</sub>* in pRG203MX | *HIS3* | This study |
| pAA18 | *lexO*-CAS9-ER-NLS-FLAG-*T<sub>ADH1</sub>* *P<sub>ACT1</sub>*-LEXA-ER-B112-*T<sub>CYC1</sub>* in pRG203MX | *HIS3* | This study |
| pAA19 | *lexO*-CAS9-NLS-FLAG-*T<sub>ADH1</sub>* *P<sub>ACT1</sub>*-LEXA-ER-B112-*T<sub>CYC1</sub>* in pRG203MX (CAS9 and LEXA-ER-B112 tail to tail) | *HIS3* | This study |
| pAA20 | *lexO*-CAS9-ER-NLS-FLAG-*T<sub>ADH1</sub>* *P<sub>ACT1</sub>*-LEXA-ER-B112-*T<sub>CYC1</sub>* in pRG203MX (CAS9 and LEXA-ER-B112 tail to tail) | *HIS3* | This study |
| pAA21 | *P<sub>tRNA-LYS</sub>*-HDV_ribozyme-sgRNA48-*T<sub>SNR52</sub>* in pRG205MX | *LEU2* | This study |
| pAA22 | *P<sub>tRNA-LYS</sub>*-HDV_ribozyme-sgRNA160-*T<sub>SNR52</sub>* in pRG205MX | *LEU2* | This study |
| pAA23 | *P<sub>GAL1</sub>*-CAS9-NLS with *P<sub>tRNA-LYS</sub>*-HDV_ribozyme-sgIR-gRNA_ mut2-*T<sub>SNR52</sub>* | *Kan* | This study |
| pAA24 | *P<sub>GAL1</sub>*-CAS9-NLS with *P<sub>tRNA-LYS</sub>*-HDV_ribozyme-sgRNApol3-01-*T<sub>SNR52</sub>* | *Kan* | This study |

